# Systematic Chromatin Architecture Analysis in *Xenopus tropicalis* Reveals Conserved Three-Dimensional Folding Principles of Vertebrate Genomes

**DOI:** 10.1101/2020.04.02.021378

**Authors:** Longjian Niu, Wei Shen, Zhaoying Shi, Na He, Jing Wan, Jialei Sun, Yuedong Zhang, Yingzhang Huang, Wenjing Wang, Chao Fang, Jiashuo Li, Piaopiao Zheng, Edwin Cheung, Yonglong Chen, Li Li, Chunhui Hou

## Abstract

Metazoan genomes are folded into 3D structures in interphase nuclei. However, the molecular mechanism remains unknown. Here, we show that topologically associating domains (TADs) form in two waves during *Xenopus tropicalis* embryogenesis, first at zygotic genome activation and then as the expression of CTCF and Rad21 is elevated. We also found TAD structures continually change for at least three times during development. Surprisingly, the directionality index is preferentially stronger on one side of TADs where orientation-biased CTCF and Rad21 binding are observed, a conserved pattern that is found in human cells as well. Depletion analysis revealed CTCF, Rad21, and RPB1, a component of RNAPII, are required for the establishment of TADs. Overall, our work shows that *Xenopus* is a powerful model for chromosome architecture analysis. Furthermore, our findings indicate that cohesin-mediated extrusion may anchor at orientation-biased CTCF binding sites, supporting a CTCF-anchored extrusion model as the mechanism for TAD establishment.

## INTRODUCTION

Interphase chromosomes are partitioned into topologically associating domains (TADs)^1–4^ which segregate into compartments of active or repressive chromatins^5–7^. TAD structures are relatively stable and resilient to environmental perturbations^8, 9^. Chromosome architecture at the TAD level is also evolutionarily conserved in eukaryotic species^4, 10, 11^. Disruption of TAD borders leads to developmental disorders and even tumorigenesis, thus underlining the importance of 3D genome organization in gene regulation^12–15^.

The establishment of chromatin architecture during embryogenesis provides an initial spatial frame which may guide proper genome organization, chromatin interaction and gene regulation in following development and differentiation processes^16^. The timing of de novo TADs formation during development has been examined in *Drosophila*, mouse, zebrafish, and human^17–21^. TAD structures form at zygotic genome activation (ZGA) and continually consolidate during early embryo development in fruit fly, mouse and human^17, 19–21^. However, in zebrafish, TADs already exist before ZGA and are lost after ZGA before being reestablished in later developmental stages^18^. This difference raises the question if the process of TAD formation is evolutionarily conserved.

Cohesin complex-mediated DNA loop extrusion was recently reported in several in vitro studies^22, 23^ and proposed as a functional mechanism underlying TAD establishment^24–26^. In cultured cells, the deletion of cohesin Rad21 alone is enough to abolish the establishment of TADs^27^. In addition, CTCF, WAPL, and PDS5 proteins participate in the regulation of TAD and loop structure formation^28, 29^. The loss of CTCF disrupts TAD insulation but not the higher-order genomic compartmentalization^30^. Rad21 and CTCF are required for TAD formation during mouse^31^ and human embryogenesis^32^, respectively, which suggests that TAD formation in cultured and embryonic nuclei is conserved and may require both factors. However, whether the mechanism of TAD formation is through cohesin-mediated extrusion initiated at cohesin loading sites and stopped upon encountering with convergent CTCF binding sites^11, 33, 34^ during embryogenesis is unclear.

Another intriguing observation in these model organisms is that transcription appears to be dispensable for TAD formation at ZGA in fruit fly and mouse^19–21^ but not in humans^32^. Moreover, a study showed that transcription could disrupt TAD borders by displacing cohesin and CTCF^35^, while others found transcription drives the formation of domain borders^36^. These opposing findings suggest that the role of transcription in TAD formation can be context-dependent or regulated by undefined factors. Thus, the mechanism underlying de novo chromosome architecture formation is still not completely understood in any of the animal models examined so far.

Before ZGA, *Xenopus* embryos undergo 12 synchronous cell cycles without gap phases^37^. Major ZGA occurs after the 12^th^ cell cycle at stage 9 when S and Gap phases appear, and interphase lengthens^38, 39^. Before major ZGA, most of the zygotic genome is transcriptionally silent. Inhibition of new translation of target proteins can, therefore, be achieved easily by injecting morpholinos into zygotes, with maternally inherited proteins maintained. This method allows us to examine how chromatin folds progressively when the amount of target protein increases during development, and thereby assess the role of specific factors in the de novo establishment of chromatin architecture which cannot be tested easily in other animal models.

To determine the evolutionary conservation and to explore the mechanism of TAD formation during embryogenesis, we examined chromosome conformation change across multiple developmental stages in wild type (WT) embryos of *Xenopus tropicalis (X. tropicalis)* and embryos in which RNAPII, CTCF and cohesin Rad21 translation have been inhibited by morpholinos either separately or in combination. By generating in situ Hi-C maps for embryos at multiple developmental stages around ZGA, our work revealed a conserved timing of TAD formation at ZGA in *Xenopus*, similar to fruit fly, mouse, and human, but not in zebrafish. In contrast to other animal models, TADs form in two waves during *Xenopus* embryogenesis, first at ZGA and then hours after ZGA, the second wave of TADs formation coincides with the elevation in CTCF and Rad21 protein concentration. Individual inhibition of CTCF and Rad21 translation weakened TAD boundaries, and inhibition of both nearly completely abolished TAD structures, suggesting an additive or synergistic contribution of CTCF and Rad21 to TAD formation. Knock-down of RNAPII also weakened TAD structures at ZGA but not for later developmental stages, implying transcription by RNAPII is required for at least the first wave of TAD formation. Interestingly, the chromatin interaction directionality was almost always stronger on one side of the TAD border instead of being roughly equal on both border sides. Accompanying with this bias, we also found enrichment of orientation–biased CTCF binding at the same TAD border side. The distribution of Rad21 showed a similar bias in enrichment. Finally, we showed human cells have these same patterns, which together indicate that a CTCF-anchored orientation-biased cohesin-mediated extrusion could be the underlying mechanism of TAD formation. In summary, besides offering evidence supporting that TAD formation during early embryo development requires RNAPII, CTCF and Rad21, our results provide a high-quality *Xenopus tropicalis* reference genome and valuable resources for further exploration of the mechanisms of genome 3D architecture establishment.

## RESULTS

### De novo Assembly of *Xenopus tropicalis* Genome Assisted by Hi-C and Single-molecule Sequencing

To determine when TADs form during *Xenopus* embryogenesis and if the process is similar in zebrafish^18^ or fruit fly, mouse, and human^19–21, 32^, we carried out Hi-C on stage 8 (s8) *X. tropicalis* embryos (before major ZGA). Unexpectedly, chromatin interaction heatmaps plotted at 100kb resolution using the previously released reference genome v9.1^40^ showed reversions, misplacements, and gaps in nearly every chromosome (Figure 1A and S1A). Therefore, to accurately characterize genome folding patterns in *X. tropicalis*, we conducted de novo genome assembly of *X. tropicalis* using Hi-C and single-molecule sequencing^41–43^ (Figure 1B). Misplacements, reversions, and gaps were mostly fixed (Figure 1C and S1B), as shown by Hi-C heatmaps (Figure 1D and S1C). The newly assembled genome is longer and with fewer gaps (Figure 1E, S1B, and Table 1). Centromere interactions can also be detected using the newly assembled reference genome (Figure S1D and S1E). We used this version of the reference genome for all of the following analyses.

**Figure 1.**
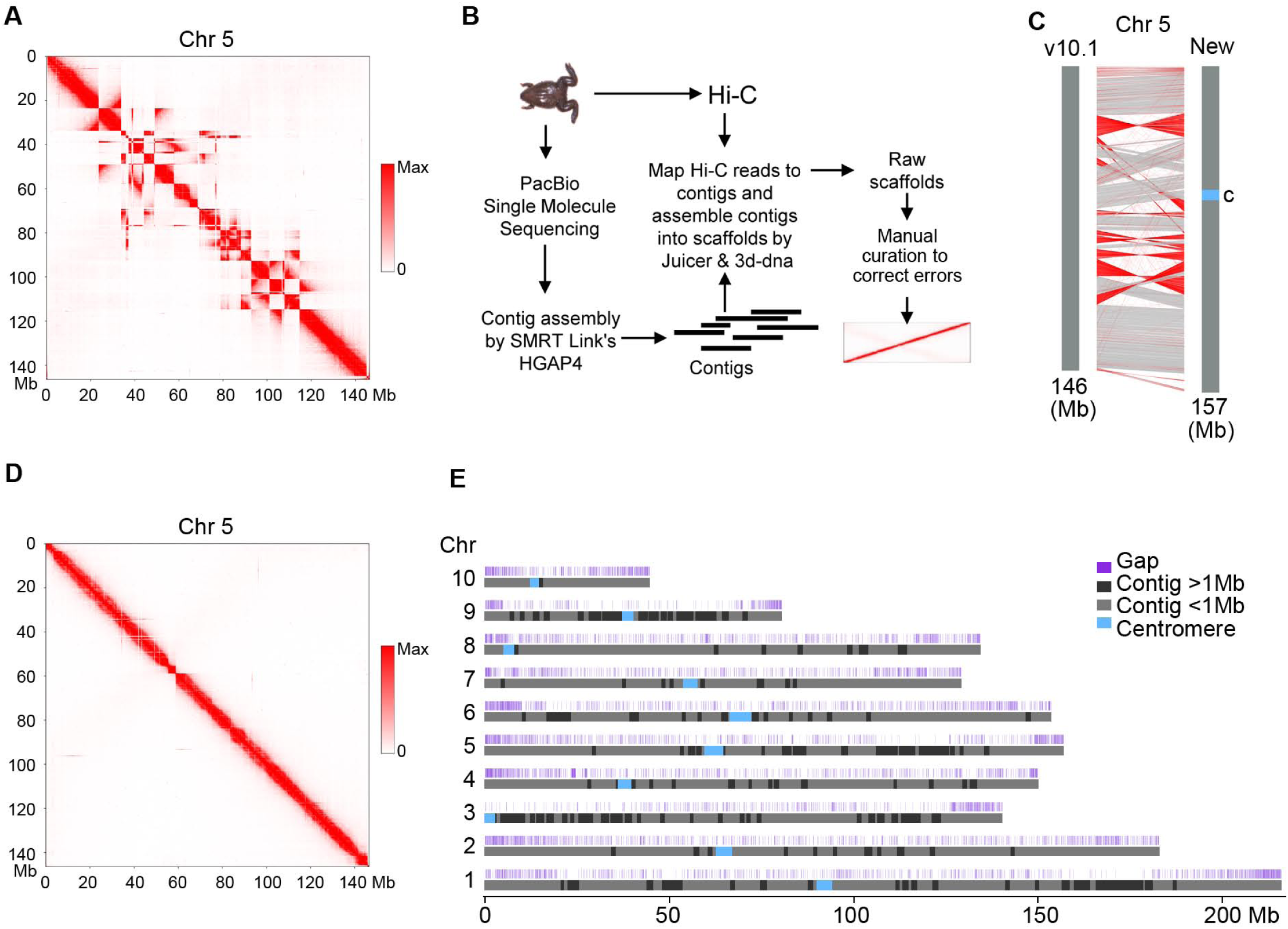
De Novo Assembly of the Reference Genome of *X. tropicalis* Assisted by Hi-C and Single-Molecule Sequencing. (A) Heatmap of chromosome 5 as an example to show assembling errors using the v9.1 reference genome of *X. tropicalis*. (B) Procedures of de novo assembly of the reference genome of *X. tropicalis*. (C) Comparison between the v9.1 and the de novo assembled chromosome 5. Red lines show sequences with orientation reversed. (D) Heatmap of chromosome 5 to show assembling errors are mostly corrected in the new version of the reference genome. (E) Ideograms of *X. tropicalis* new reference pseudomolecules. The top track shows positions of gaps (dark blue). Contigs longer than 1Mb are shown in black, and contigs shorter than 1Mb are shown in light grey. See also Figure S1.

**TABLE 1.** De Novo Assembly Statistics for the Xenopus tropicalis

|  | *Xenopus tropicalis* New version | *Xenopus tropicalis* v9.1 |
| --- | --- | --- |
| Total contig length (bp) | 1,481,767,000 | 1,369,865,000 |
| Number of contigs | 9,633 | 61,330 |
| Contig N50 | 377,941 | 71,041 |
| Contig N90 | 59,908 | 11,781 |
| Max contig length | 7,388,000 | 671,191 |
| Gap | 3,839,000 | 70,533,000 |
| Total scaffold length (bp) | 1,485,606,000 | 1,440,398,000 |
| Number of scaffolds | 1,956 | 6,822 |
| Scaffold N50 | 150,158,000 | 135,135,000 |
| Scaffold N90 | 80,475,000 | 1,477,000 |

### Datasets Description

We used PacBio single-molecule sequencing (Table S1) and Hi-C for de novo genome assembly. We generated 21 high-quality Hi-C datasets. At least two biological replicate libraries, unless otherwise stated, were generated and sequenced (Table S2). Genome-wide resolution for most Hi-Cs was below 10kb calculated as described ^11^. We generated 17 ChIP-seq datasets using CTCF, Rad21, and RNAPII antibodies on wild type embryos at stages 9, 11, and 13, and morpholino injected embryos at stages 9 and 13. We also generated ChIP-seq datasets for CTCF and Rad21 on brain and liver cells (Key Resources Table).

### TAD Structures Appear at the Onset of Major ZGA in *Xenopus tropicalis*

The timing of de novo establishment of TAD structures varies during early embryo development among evolutionarily distant animals ^18–21, 32^. This difference prompted us to examine when the 3D chromatin architecture is established in *X. tropicalis*, an important model for developmental and genetic research, which is evolutionarily close to fish but more distant from insects and mammals.

We first generated in situ Hi-C maps on hand-picked non-mitotic (S-phase) s8 embryos, which are at minor ZGA 5 hours post-fertilization (hpf) (Figure 2A). Visual inspection of high resolution (5kb) chromatin contact heatmaps failed to reveal any distinct patterns, indicating the lack of structural organization before major ZGA (Figure 2B). Synchronized and rapid cell cycle division before major ZGA may, therefore, prevent the establishment of stable chromatin structures in *X. tropicalis*. To determine if chromatin structures emerge when rapid synchronized cell division ends, we carried out in situ Hi-C on stage 9 (s9) embryos when major ZGA begins. Though weak, TAD-like structures appeared across chromatin contact heatmaps (Figure 2B), suggesting that TAD structures start to form at the major ZGA stage in *Xenopus,* similar to fruit fly, mouse, and human^19–21, 32^ but not zebrafish^18^.

**Figure 2.**
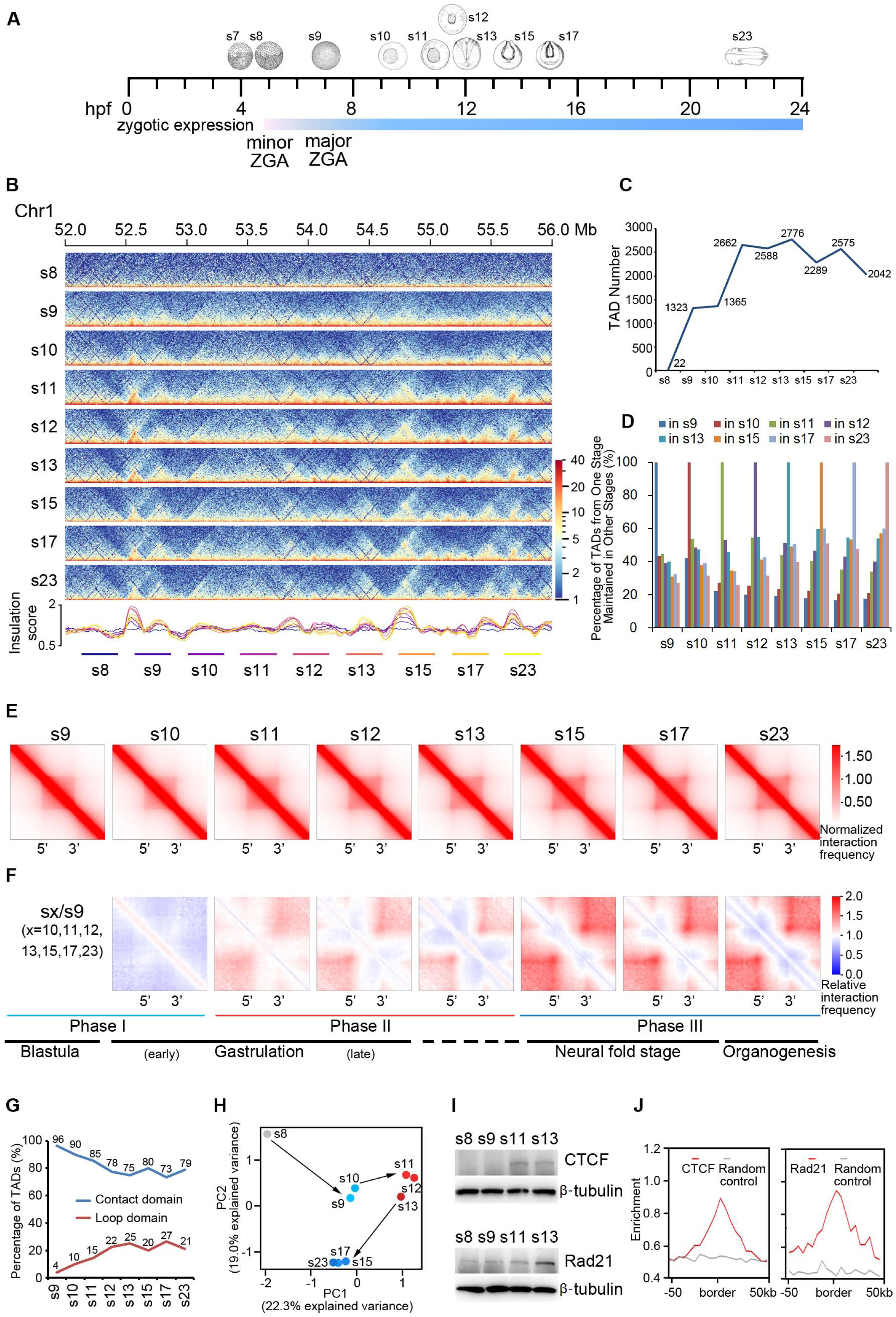
Two-wave and Three-Phase TAD Establishment in *X. tropicalis*. (A) Schematic representation of the ten developmental stages(s) examined by in situ Hi-C. (B) Chromatin interaction frequency mapped at 5kb resolution. (C) Number of TADs identified at the eight developmental stages. (D) Overlapping of TADs between different stages. (E) Heatmaps of aggregated TADs for the eight developmental stages. (F) Interaction frequency of aggregated TADs from s10 to s23 was normalized against s9. Three phases of change in TAD structure are shown below with the developmental stages also shown. (G)Percentage of contact and loop domains at the eight developmental stages. (H) Principle component analysis of insulation scores of Hi-Cs for the nine developmental stages. (I) Western blotting of CTCF and Rad21 for embryos at four developmental stages. (J) Enrichment of CTCF and Rad21 at TAD borders in embryos of stage 9. See also Figure S2.

### Two-Wave and Three-Phase TAD Formation during Embryogenesis in *Xenopus tropicalis*

Next, we examined changes in chromatin conformation at later developmental stages (stages 10, 11, 12, 13, 15, 17 and 23, abbreviated as s10, s11, s12, s13, s15, s17 and s23, respectively) after major ZGA (Figure 2B). In contrast to fruit fly in which TAD structures are mainly established at major ZGA, TAD boundaries in *Xenopus* increased from approximately 1300 at s9 and s10 to 2662 at s11. This level was maintained throughout later developmental stages (Figure 2C) with relatively stable median TAD sizes (Figure S2A). Consistent with this pattern, the percentage of the genome that is folded into TADs positively correlated with the number of TADs established at each stage (Figure S2B), suggesting there are two major TAD establishment stages which are separated between s10 and s11. TADs formed overlap more with adjacent developmental stages (Figure 2D). Taken together, these results support a two-wave TAD establishment pattern during *X. tropicalis* embryogenesis after major ZGA. This finding is similar to fruit fly in which active long-range chromatin loop formation also occurs in two waves during early embryogenesis ^17^.

To compare if TADs formed at different stages are similar or different in structure, we aligned all domains and calculated the average interaction frequency within and between TADs. For domains at s9 and s10, the interaction loops formed between borders were not apparent, suggesting domains at these two stages are mostly contact domains (Figure 2E). Chromatin interaction frequency between borders appeared at s11 and became increasingly stronger in later stages (Figure 2E), indicating the establishment of loop domains.

We examined differences in chromatin interactions further by normalizing the chromatin interaction frequency of aggregated TADs against s9. Consistent with the previous observation, TADs formed at s9 and s10 were similar in chromatin interaction frequency (Figure 2F). Surprisingly, TADs formed at s11, s12, and s13 were similar to each other, as were TADs formed at s15, s17, and s23 (Figure 2F). These results indicate that TADs formed in the two waves were not only separated between s10 and s11 but in fact, their structure can be further divided between s13 and s15. Since we did not conduct Hi-C for stage 14 (s14) embryos, we could not rule out the exact timing of the difference between s13 and s14 or between s14 and s15. Interestingly, the three-phase changes in TAD structures almost correspond to the four consecutive developmental stages of *X. tropicalis* (Figure 2F).

Next, we counted the number of loop domains and contact domains and calculated the percentage of these two types of domains. Our results show that the percentage of loop domains steadily increased from 4% at s9 to 15% at s11 and remained stably above 20% from s12 to the later stages (Figure 2G). This finding suggests that loop domain establishment is relatively slower than overall TAD formation. PCA analysis on the directionality index separated s9 and s10 into one group, s11, s12, and s13 into a second group, and s15, s17, and s23 into a third group (Figure 2H). This result reflects the significant changes occurring at several distinct transition points between s8 and s9, s10 and s11, and s13 and s15 in the chromatin interaction pattern of the *X. tropicalis* genome. However, whether these changes are functionally significant, are not determined at this time.

### Two-Wave TAD Formation Correlates with CTCF and Rad21 Expression

In vertebrate genomes, CTCF motifs at TAD borders are, in general, convergently paired^11, 33, 34^. In line with previous studies, our analysis also revealed similar convergent CTCF motif orientation for TADs identified at the different stages of development in *X. tropicalis* (Figure S2C). This result suggests that TAD formation in *X. tropicalis* requires CTCF as well. To explore this further, we examined changes in protein expression for CTCF and Rad21 by western blotting. CTCF was barely detected at s8 and appeared slightly at s9 (Figure 2I), which accompanies the first-wave of TAD formation (Figure 2C). CTCF protein levels increased dramatically at s11 (Figure 2I), which corresponds to the start of the second-wave of TAD formation (Figure 2C). In contrast, Rad21 was detected at both s8 and s9 and increased at s11 (Figure 2I), suggesting that cohesin alone is not enough for the de novo establishment of the first-wave of TADs. ChIP-seq analysis showed that both CTCF and Rad21 are already bound to DNA at s9 and remained bound at s11 (Figure 2J and S2D). Taken together, these results demonstrate that the two-wave TAD formation is closely linked to the increase in CTCF and Rad21 protein concentration, further implying the requirement of both CTCF and cohesin Rad21 for TAD establishment.

### Directionality Index Is Preferentially Higher at One Side of TAD Borders

Chromatin at TAD borders preferentially interacts with fragments inside TADs. This bias in chromatin interaction can be measured as the directionality index (DI)^4^. To explore the underlying cause of the directionality, we aligned TADs at 5’ and 3’ borders and extended 5 bins (5Kb per bin) up- and down-stream of the two borders. We clustered TADs based on the DIs of each domain at the two borders. Surprisingly, we found that TADs from s13 are grouped into three distinct clusters (Figure 3A and S3A). Absolute DI values upstream and downstream of the borders in clusters 1 and 3 were strikingly higher at one side of the border (Figure 3A). Most TADs were in cluster 2 with similar average absolute DI values at both borders (Figure 3A). We also observed similar enrichment patterns for CTCF and Rad21 binding across the three clusters (Figure 3B and 3C). We further divided TADs in cluster 2 into five sub-clusters of an equal number of TADs (Figure S3A). We calculated the directionality bias index for each cluster and found absolute DI values were also bigger at either the 5’ or the 3’ border (Figure 3D). These results together strongly suggest that the difference in the DI patterns across borders could be caused by the orientation-biased and enrichment-biased binding of both CTCF and cohesins at one side of the TAD border. Examples of TAD for each cluster are shown in Figure 3E.

**Figure 3.**
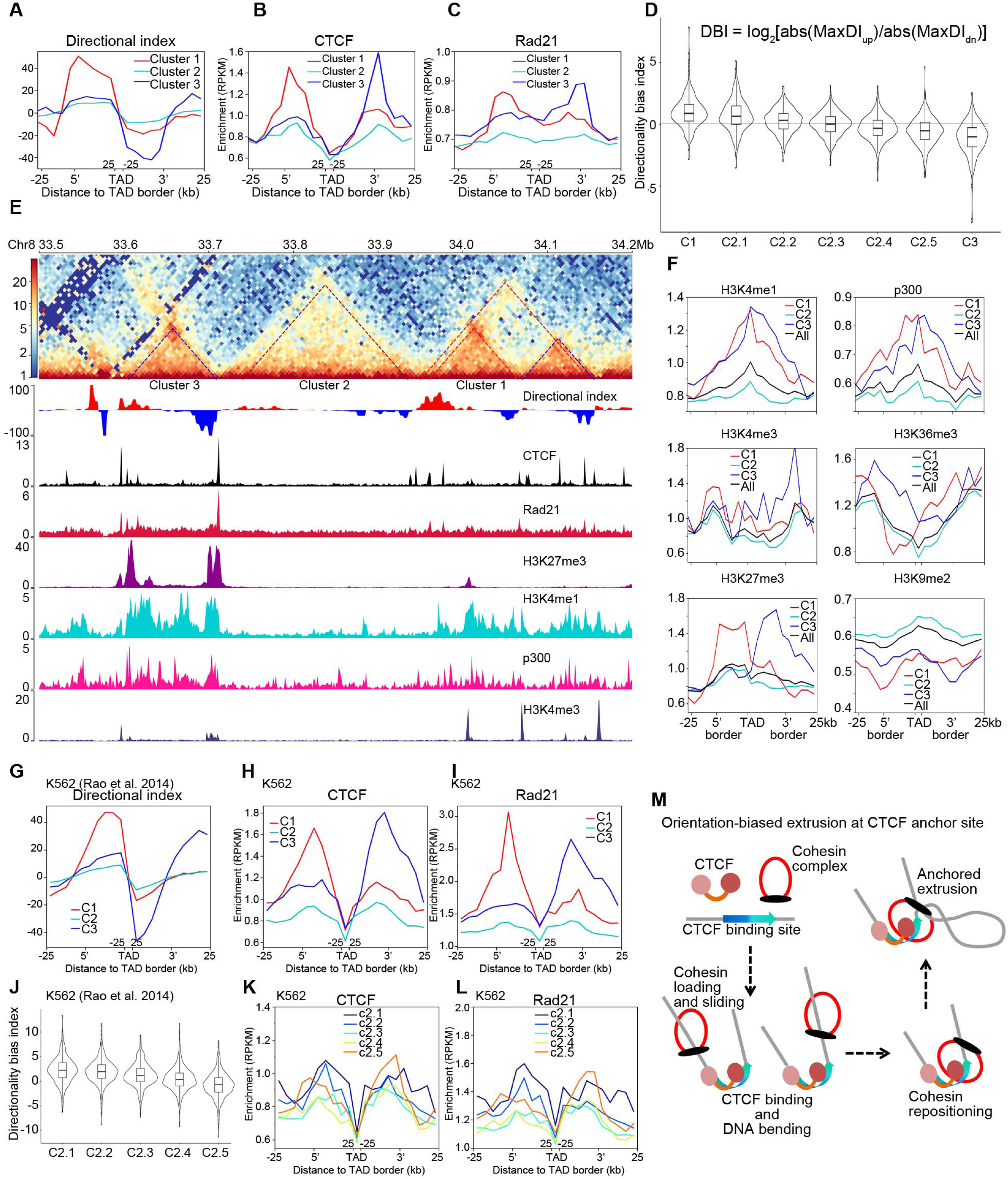
Orientation-Biased CTCF and Rad21 Enrichment at TAD Borders of Higher Directionality Index Values. (A) Directionality index for three clusters of TADs identified in embryos at stage 13; 344, 2035 and 397 TADs in Cluster 1, 2 and 3, respectively. (B) CTCF enrichment is biased to borders with higher directionality index values. (C) Rad21 enrichment is biased to borders with higher directionality index values and drops toward the border on the other side of TAD. (D) Directionality bias index (DBI) for TADs of cluster 1, five sub-clusters of clusters 2, and cluster 3. According to the rank of directionality index strength, TADs in cluster 2 are divided into five sub-clusters. (E) Examples of TADs for the three different clusters. (F) Histone modifications and p300 enrichment patterns across the borders of three clusters of TADs. (G)Directionality index across borders of the three clusters of TADs in K562 cells. (H) Enrichment of CTCF is biased to borders with higher directionality index values for the three clusters of TADs. (I) Enrichment of Rad21 is biased to borders with higher directionality index values for the three clusters of TADs. (J) Directionality bias index for the five sub-clusters of cluster 2. According to the rank of directionality index strength, TADs in cluster 2 are divided into five sub-clusters. (K) Enrichment of CTCF is biased to borders with higher directionality index for the five sub-clusters of cluster 2. According to the rank of directionality index strength, TADs in cluster 2 are divided into five sub-clusters. (L) Enrichment of Rad21 is biased at borders with higher directionality index for five cluster 2 sub-clusters. According to the rank of directionality index strength, TADs in cluster 2 are divided into five sub-clusters. (M) Model of Cohesin-mediated directional extrusion anchored at orientation-biased CTCF binding site. See also Figure S3.

The unexpected revelation of DI bias at TAD borders shows that simple aggregation analysis of all TADs conceals rich structural information, which may help us to understand the genome folding mechanism. Indeed, the analysis of all TADs again showed undistinguishable DI patterns for CTCF and Rad21 enrichment at TAD borders (Figure S3D, S3E, and S3F).

Similarly, the enrichment patterns of H3K4me1, p300, H3K4me3, H3K36me3, H3K27me3, and H3K9me2 at TAD borders were different for each cluster (Figure 3F). Whether these epigenetic modifications are important for TADs establishment is not determined at this time. However, it is reasonable to speculate that chromatin states can affect the process of cohesin-mediated extrusion as for super-enhancers^44^. Surprisingly, neither the density of genes nor RNA expression was found enriched at the borders of either cluster of TADs (data not shown).

To find out if the biased pattern of DI and enrichment of CTCF and Rad21 are conserved through evolution, we carried out similar analysis using previously published data for human K562^11^ and *Drosophila* S2 cells^45^. Strikingly, we found even more obvious patterns in human K562 for all the three parameters analyzed (Figure 3G, 3H, 3I and S3G, S3H, S3I) and even the five sub-clusters of cluster 2 showed obvious bias in DI, CTCF, and Rad21 binding (Figure 3J, 3K, and 3L). Though the DI bias was also found for TADs in S2 cells (Figure S3J and S3K), CTCF enrichment was not biased for any of the three clusters (Figure S3L), consistent with the lack of convergent CTCF motifs at TAD borders in the *Drosophila* genome^46^. Taken together, these results imply that orientation-biased CTCF binding may provide an anchor site for the cohesin complex to initiate directionality-biased extrusion in vertebrate genomes (Figure 3M), at least for *X. tropicalis* and human. However, the mechanism underlying the directionality bias at TAD borders in *Drosophila* can be different from vertebrates because of the lack of CTCF orientation-biased motif.

### CTCF and Cohesin Rad21 Are Required for De Novo TAD Formation

ChIP-seq results of CTCF and Rad21 at s9 showed global occupancy of both factors (Figure S2D and S2E) despite only a trace amount of detectable CTCF protein expression (Figure 2I). This observation suggests even low levels of CTCF are enough for DNA binding and sufficient for the first-wave of TAD formation at the beginning of ZGA. We also found that the second-wave of TAD formation is correlated with the increase of CTCF and Rad21 protein amount (Figure 2C and 2I). Recently, CTCF was shown to be critical for TAD formation during human embryogenesis^32^. Together, these results suggest a common conserved mechanism for a two-wave establishment of TADs in *Xenopus* and human embryos that requires both CTCF and Rad21.

To test this hypothesis, we knocked down CTCF and Rad21 with morpholinos targeting them individually or in combination. Morpholinos injected into fertilized eggs inhibit new translation with maternally inherited proteins maintained. This incomplete depletion of target proteins thus provides a unique opportunity to observe if changes in protein concentration affect chromatin architecture. Morpholinos targeted against CTCF and Rad21 efficiently reduced both proteins to background levels at s13 (Figure 4A-C). The reduction in CTCF or Rad21 expression resulted in a minor decrease in overall CTCF and Rad21 binding to chromatin (Figure S4A), suggesting that background levels of CTCF and Rad21 can still support DNA binding. However, knock-down of either factor considerably weakened TAD structures. The Arrowhead corner scores for TADs were reduced after knock-down of both factors (Figure 4D). We also calculated the insulation scores for borders before and after knock-down and found that insulation at most borders dropped, especially after the combined knock-down of CTCF and Rad21 (Figure S4B), but not in embryos treated with control morpholino (data not shown). And TAD structures were almost completely abolished with the simultaneous reduction of both CTCF and Rad21 (Figure 4E and S4C). Knock-down of CTCF, Rad21, or both significantly reduced the number of TADs (Figure 4F) without affecting the median size of the TADs (Figure S4C) and the percentage of genome folded into TADs was proportional to the number of TADs (Figure S4D). Similar effects of CTCF and Rad21 on TAD structures were also observed in s9 embryos (Figure S4E). Taken together, these results strongly support that both CTCF and Rad21 are required for the de novo establishment of TADs during *X. tropicalis* embryogenesis. The prerequisite of both factors is likely to be conserved for TAD formation independent of developmental stages and cell types of vertebrate animals

**Figure 4.**
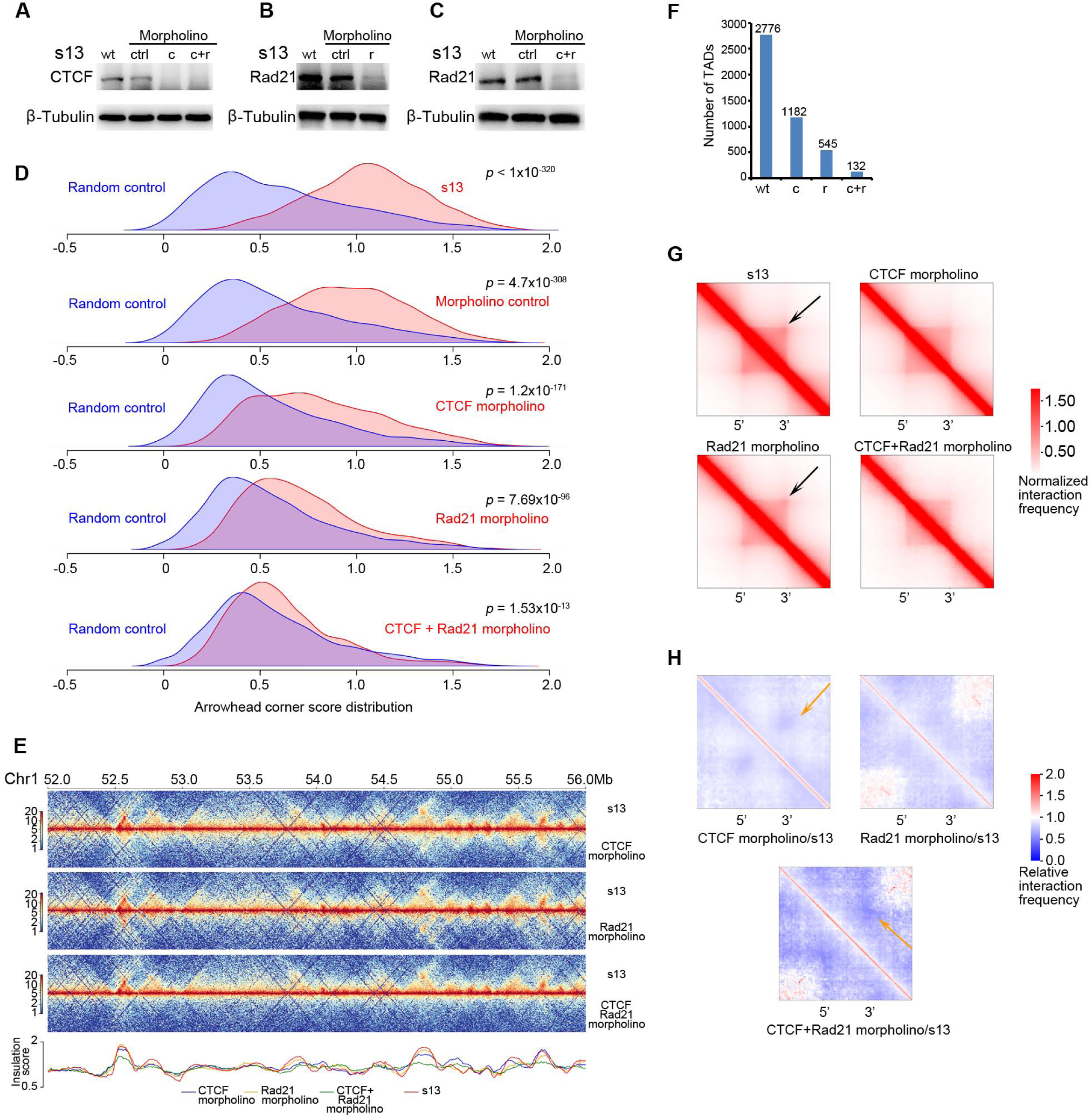
Requirement of CTCF and Cohesin Rad21 for TAD Establishment in *X. tropicalis* Embryos. (A), (B) and (C) Western blotting of CTCF and Rad21 knock-down by morpholinos in embryos at stage 13. Wild type (wt), control (ctrl), CTCF morpholino (c), Rad21 morpholino (r). (D) Arrowhead corner score distribution for wild type, morpholino control, and knock-downs with CTCF morpholino, Rad21 morpholino and combined CTCF and Rad21 morpholinos. All experiments were carried out on stage 13 embryos (*P* values, calculated using Mann-Whitney U Test). (E) Example region to showing the knock-down effect on TAD structures. (F) Number of TAD after knock-down. (G)Heatmaps of aggregated TADs. Black arrows point to interacting borders. (H) Interaction frequency of aggregated TADs normalized against wild type s13. Orange arrows point to weakened border interaction. See also Figure S4.

Interactions between TAD borders become apparent at s13 in *X. tropicalis* (Figure 2E and 2F). Knock-down of CTCF not only compromised TAD formation overall, but it also weakened the interactions between TAD borders (Figure 4G and 4H). In contrast, knock-down of Rad21 severely weakened TAD structures but not interactions between TAD borders (Figure 4G and 4H). As expected, the combinational knock-down of CTCF and Rad21 abolished both TADs and loop interactions between TAD borders (Figure 4G and 4H). Taken together, these results indicate that the roles of CTCF and Rad21 in TAD establishment are different. CTCF appears to contribute more to loop formation^47, 48^, while cohesin Rad21 seems to contribute more to domain structure.

### RPB1, a Component of RNAPII, Plays an Important Role in the First Wave of TAD Formation and Embryo Development

The role of transcription in TAD formation is controversial. However, a recent study of human embryogenesis supports that transcription is important for TAD establishment. RNA transcription is carried out by the RNAPII complex, in which the protein RPB1 is a critical component. RPB1 is expressed before ZGA and remains at a similar level until dramatically increasing at s11 (Figure 5A). To determine whether RNAPII is important for TAD establishment during *X. tropicalis* embryogenesis, we inhibited RNAPII activity by depleting RBP1 protein levels. Morpholinos against RPB1 efficiently reduced the protein to background levels in embryos that developed to s10 (Figure 5B). Except for a few hundreds of genes, transcription of most genes was not significantly affected by RPB1 depletion (data not shown), a result is likely due to the existence of maternally inherited proteins.

**Figure 5.**
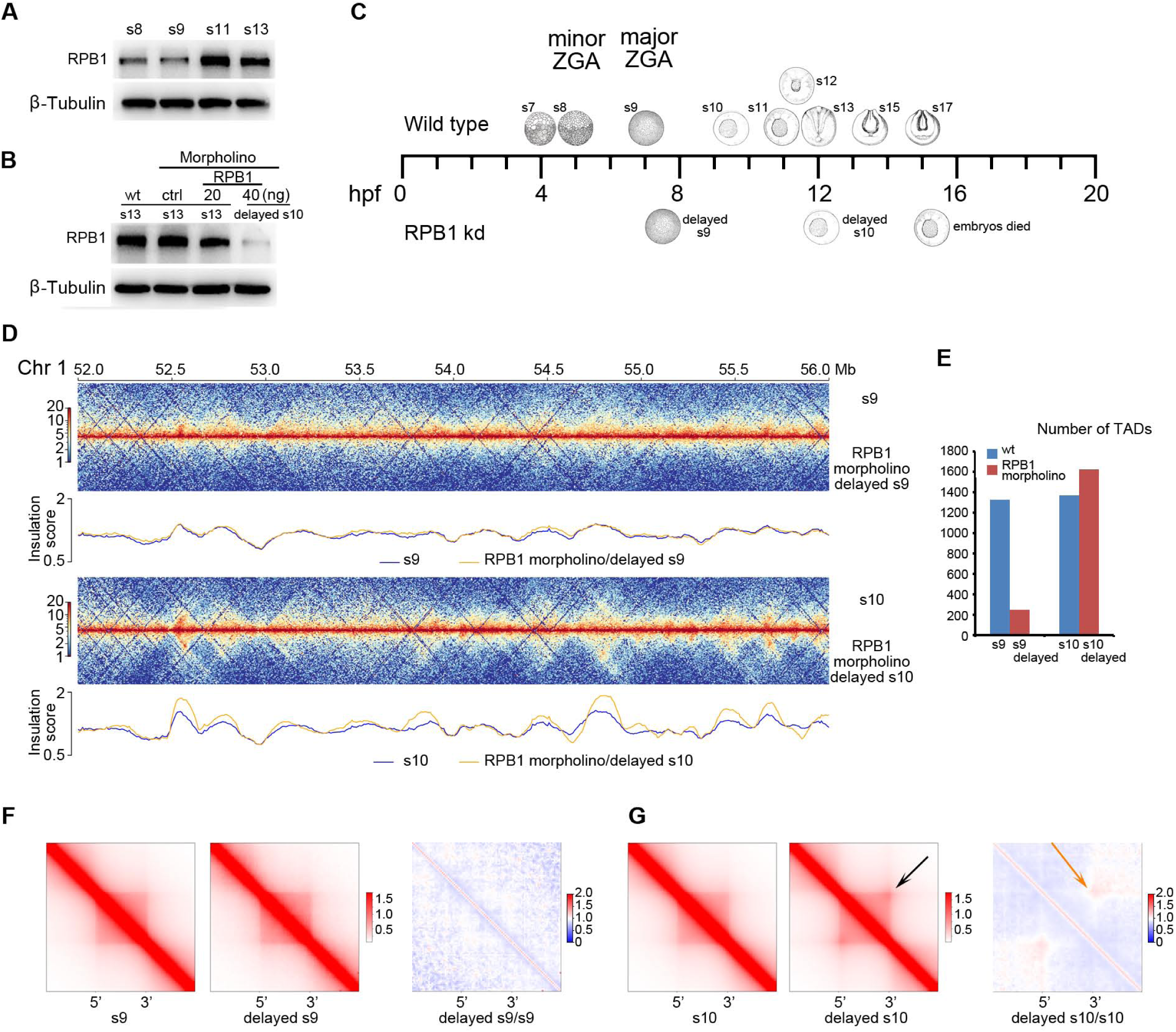
RPB1 Knock-Down Effects on TAD Establishment. (A) Western blotting of RPB1 in wild type embryos at the four developmental stages. (B) Western blotting of RPB1 knocked-down with 20 or 40ng of morpholinos. (C) Schematic representation of stages s9 (delayed) and s10 (delayed) caused by RPB1 knock-down. (D) Example of a region to show the knock-down effect on TAD structures. (E) Number of TADs. (F) Aggregated TAD analysis for embryos of wild type s9 and RPB1 knock-down delayed s9, and comparison of them by dividing delayed s9 over wild type s9. (G)Aggregated TAD analysis for embryos of wild type s9 and RPB1 knock-down delayed s10, and comparison of them by dividing delayed s10 over wild type s10. See also Figure S5.

We first examined the effect of RNAPII inhibition during embryogenesis. RPB1 knock-down dramatically delayed embryo development, with s9 taking about half an hour more than usual, while s10 took nearly five and a half hours longer, about the same time for wild type embryos to reach s13. RPB1 knock-down embryos eventually die before reaching s11 (Figure 5C). These results show that RNAPII is very important for early embryo development.

Next, we examined if RNAPII is important for TAD establishment during *X. tropicalis* embryogenesis. TADs that were formed at delayed s9 were similar to the ones formed at normal s9 (Figure 5D), and interactions between TAD borders were not observed (Figure 5F). Interestingly, in embryos at delayed s10, loop interactions between TAD borders were established, which were not formed in WT embryos at s10 (Figure 5G). Surprisingly, the depletion of RBP1 did not affect TAD structures on delayed s10 embryos (Figure 5D, 5E, S5A, S5B, and S5C), which is different from CTCF/Rad21 knock-down, whose effects on TAD formation is persistent through early developmental stages, at least until s13. In contrast, knock-down of RPB1 dramatically weakened the TAD structure at s9 delayed embryos (Figure 5D, 5E, S5D, S5E, and S5F).

These results together indicate that the first-wave of TAD establishment is more sensitive to the reduction of transcription. However, whether this is caused directly by compromised transcription by RNAPII or by indirect effects from RNAPII reduction remains to be determined. Without newly synthesized RPB1, each division halved the amount of RNAPII in each cell. Because of this, we speculate that the background amount of RNAPII provides just enough protein for TAD establishment in delayed s10 embryos.

### Progressive Genome Compartmentalization after ZGA

Separation of chromatin into active and repressive compartments A and B is another prominent structural feature of metazoan genomes. Embryogenesis provides a unique window to observe if the process of chromosome compartmentalization and TAD establishment are linked or uncoupled^30^. We examined compartmentalization by plotting chromatin contact heatmaps at 100kb resolution. Visual inspection of heatmaps revealed continuous expansion of long-range chromatin interactions from s8 to s23 with the appearance of compartment-like patterns starting as early as s13 (Figure 6A and 6B). A zoomed-in view of two 30Mb regions on chromosome 2 revealed a more obvious initiation of compartmentalization beginning at s9, the same time as TADs are initially established (Figure 6C and S6A). In comparison to the two-wave TAD formation process, newly segregated compartments continuously emerge through later developmental stages reaching the highest level at stage 23 (Figure 6D). PCA analysis of eigenvector values shows that compartmentalization of the genome can be separated into five phases (Figure 6E), with s9 and s10 as the first phase of compartmentalization and TAD establishment (Figure 2F and 2G). Taken together, these results show compartments are continually refined after ZGA initiation and change progressively into new states that are independent of those in TAD formation.

**Figure 6.**
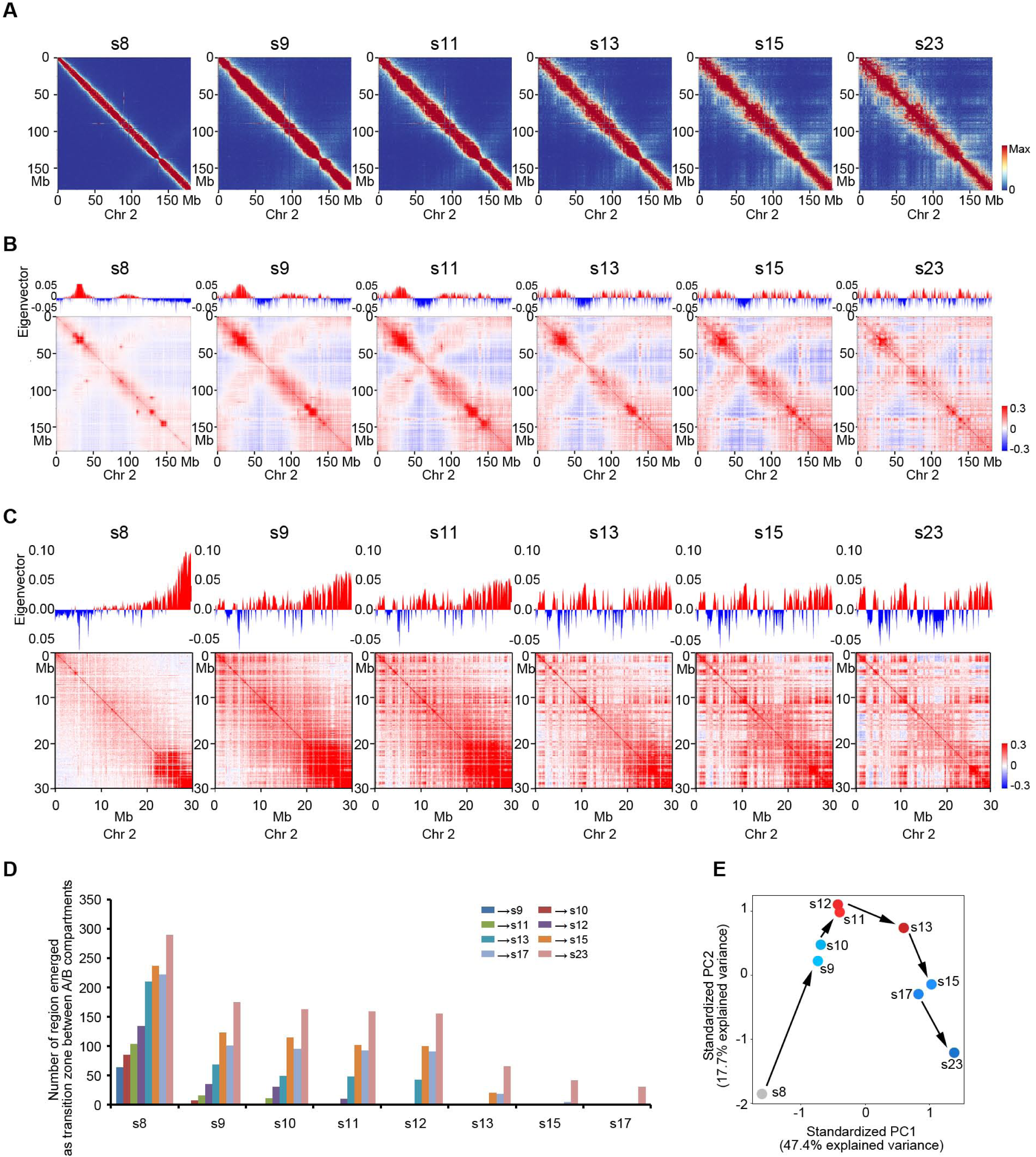
Continuous Compartmentalization during Embryogenesis. (A) Heatmaps of chromosome 2 plotted at 50kb resolution at multiple developmental stages. (B) Pearson correlation Hi-C matrices for chromosome 2 at 50kb resolution at multiple developmental stages. (C) Pearson correlation Hi-C matrices of an example region between 0-30Mb in chromosome 2. (D) Number of regions emerged as transition zone between A/B compartments compared to different start stage as indicated. (E) Principle component analysis of eigenvectors of Pearson correlation Hi-C matrices at multiple developmental stages. See also Figure S6.

### TAD and Compartment Structure Are Cell Type-Specific in Adult Frog

Chromosome 3D architecture is relatively stable in many types of mammalian cells and even conserved in different species^4, 10, 11^. Whether chromosome architecture is conserved in different cell types of *Xenopus* is unknown. To address this question, we carried out Hi-C on adult brain and liver cells of *X. tropicalis*. A comparison of chromatin interaction heatmaps for chromosome 2 shows apparent differences in interaction patterns between brain and liver cells (Figure 7A). Pearson’s correlation coefficient analysis of eigenvector further supports the compartmentalization of chromosomes is distinct between the two cell types (Figure 7B).

**Figure 7.**
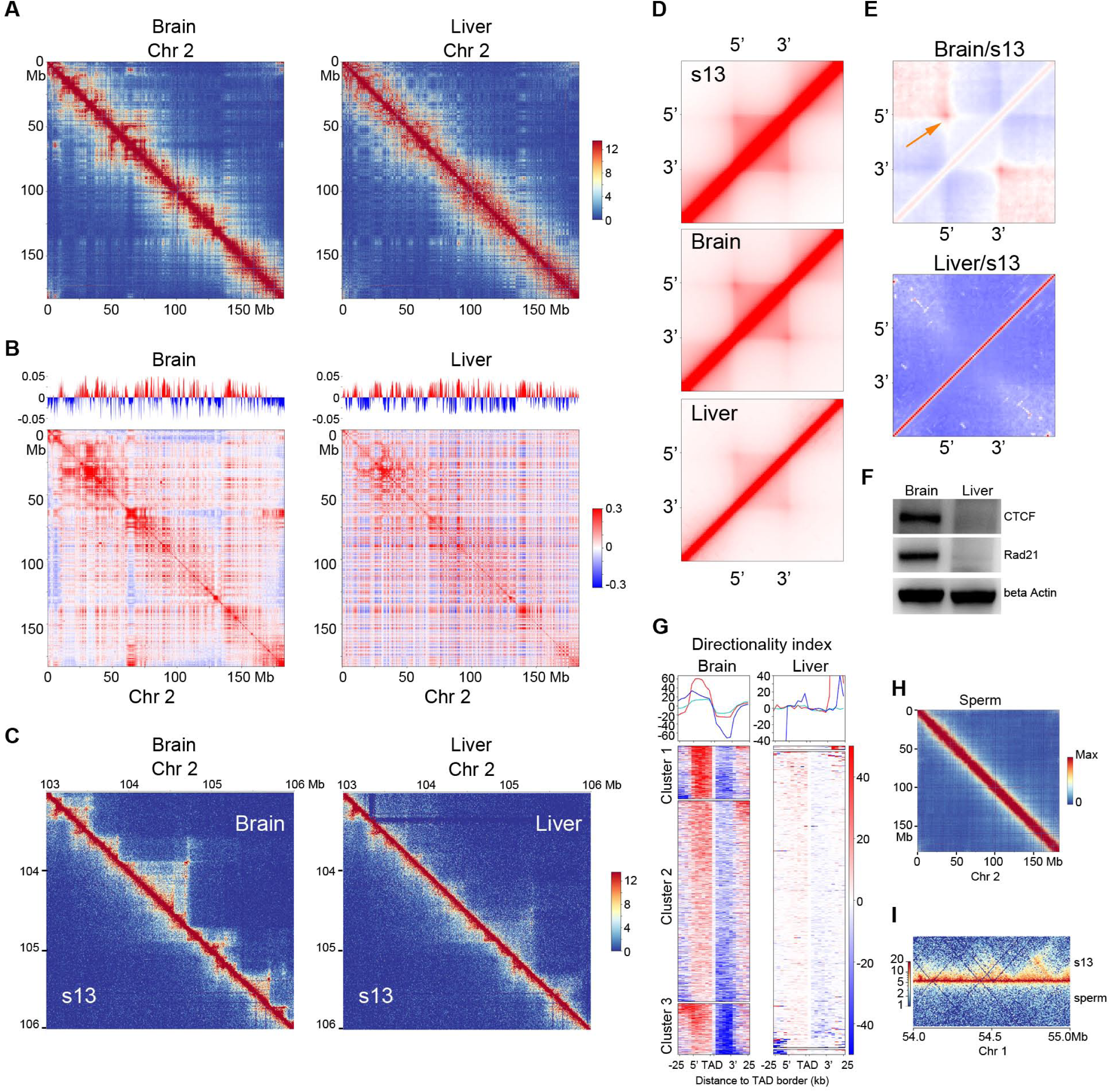
TADs Lost in Mature Liver Cells Correlates with Low Level Expression of CTCF and Rad21. (A) Heatmaps of chromosome 2 plotted at 50kb resolution for brain and liver cells. (B) Pearson correlation Hi-C matrices for chromosome 2 at 50kb resolution for brain and liver cells. (C) Heatmaps of an example region in chromosome 2 showing the gain and the loss of TAD structures in brain and liver cells comparing to s13. (D) Aggregated TADs of brain, liver, and s13 embryo cells. (E) Normalization of aggregated TADs against s13 embryos. (F) Directionality index cluster for TADs of brain and liver cells. (G)Western blotting of CTCF and Rad21 in brain and liver cells. (H) Heatmap of chromosome 2 plotted at 50kb resolution for mature sperm. (I) An example region to show the lack of TAD structure in sperm genome. See also Figure S7.

Because the establishment of TADs and compartments are uncoupled processes, we wondered if TAD structures are conserved in different cell types of *X. tropicalis*. We identified 3107 and 495 TADs in brain and liver cells, respectively (Figure S7A, S7B, and S7C). A comparison of chromatin interactions between cells from s13 with brain and liver cells revealed stark differences. TAD structures were more evident in brain cells than in s13 embryos (Figure 7C). However, TAD structures in liver cells were much weaker compared to s13 cells (Figure 7C). The distribution of Arrowhead corner scores for brain cells was consistently shifted to the right compared to liver cells (Figure S7D). Aggregation of TADs showed that interactions between TAD borders were also more evident in brain cells than in liver cells (Figure 7D). When we normalized brain and liver aggregated TADs against those from s13, we found that TADs in brain cells were more enriched with interactions between TAD borders than liver cells (Figure 7E). Together, these results suggest that the chromosome architecture is organized in a cell-type specific manner in *Xenopus*.

Our CTCF and Rad21 knock-down results in embryos showed that CTCF and Rad21 were required for TADs formation and also for loop structure formation. We speculated that the lack of TAD structures in liver cells could be due to a low concentration of CTCF and Rad21. Indeed, western blot analysis showed that both CTCF and Rad21 were highly expressed in brain cells but barely detectable in liver cells (Figure 7F). And DI clustering showed similar biases in the strength of chromatin interaction directionality at borders of TADs in brain cells in which TADs were dominant (Figure 7G), but not in the liver in which TADs were weak, suggesting directionally anchored extrusion was weak (Figure 7G). This observation further supports our hypothesis that TAD structure establishment during embryogenesis and in terminally differentiated cells are likely through conserved mechanisms.

We also examined the genome architecture of mature sperm cells from *X. tropicalis*. Compared to mouse sperm cells^49, 50^, neither compartments (Figure 7H) nor TAD structures (Figure 7H) could be detected in *X. tropicalis* sperm cells. These results together show that the genome architecture in terminally differentiated cells could be extremely different. How these different structures are established, and if they are important for cell-type specific gene expression remains to be explored.

## DISCUSSION

The establishment of chromosome architecture at the beginning of a new life cycle in a metazoan can be pivotal for guiding or dictating the chromatin organization and the precise regulation of gene transcription in a temporally and spatially controlled manner. Previous studies have shown that chromosome architectures are established at ZGA in *Drosophila*, mouse, and human^19–21, 32^ but not in zebrafish^18^. Here, our results show that in *Xenopus*, TADs are also established at ZGA, suggesting a conserved process in an evolutionally distant model organism.

In addition, our systematic analysis of chromosome architecture in multiple developmental stages allowed us to show TADs are established at two distinct developmental stages, firstly at s9 and then at s11. As the embryos develop, TADs actually continuously change their internal structure, with enhanced interactions at TAD borders. Similarly, two waves of active long-range chromatin loop formation was reported in fruit fly^17^. In other model organisms, the number of TADs also increases continuously after ZGA^20, 21^, but whether their internal structures change similar to *X. tropicalis* remains unknown. Thus, it would be interesting to determine in future work if the continuous change in TAD internal structure is another conserved process during embryo development in other model organisms.

Besides establishing two waves of TAD, our work revealed that changes in TAD internal structures further separated the two waves into additional three groups that approximately correspond to three distinct developmental phases. One possible explanation is that the composition of cell types may be similar during a specific phase, which could cover several developmental stages. Another possibility is that our Hi-Cs experiments contained a mixture of all cell types, which likely masked and averaged the differences in chromosome architecture in the different types of cells, even though there are limited cell types at early developmental stages.

Enrichment of RNA expression and high gene density are two characteristics frequently observed at TAD borders in other systems^1, 3, 4^. However, we failed to identify either of these two features at TAD borders in *X. tropicalis*, which could also be caused by the heterogeneity in the cells we used. Recent work revealed that cells could be different at as early as two- to four-cell stages during mouse embryo development^51^. At s9, at least three lineages of cells already exist in *X. tropicalis* embryos. These findings, therefore, highlight the importance of separating different types of cells for Hi-C at early developmental stages in the future.

In cultured cells, the role of CTCF and Rad21 has been intensively examined, with both proteins shown to be important for TAD establishment. There is also evidence that supports TAD establishment during embryogenesis may be through the same mechanism requiring cohesin-mediated extrusion and CTCF protein^31^. In our work, we show that knock-down of CTCF and Rad21 disrupt TAD formation but not the embryo development. In addition, we found that liver cells, which express low levels of CTCF and Rad21, have no distinct TAD structures. Together, we confirm that TAD formation requires both CTCF and cohesin, and this is likely independent of cell types and developmental stages in vertebrates.

We detected Rad21 expression by western blotting at s8 and s9, before and at ZGA. Our work suggests that inhibition of protein translation by morpholino does not eliminate preexisting target proteins in oocyte. Thus, the knock-down effects observed from our morpholino experiments are caused not by complete elimination of target protein, but instead by the under-expression of the target protein. In fact, after knock-down, CTCF and Rad21 binding are still observed across the genome. However, TAD formation was severely disrupted. Together, these findings indicate that protein concentration is another factor, which should be considered when studying chromatin organization. In support of this, studies on phase separation showed that the density of transcription factor binding to DNA motif affects higher-order chromatin structure formation^52, 53^.

Another intriguing question related to the mechanism of TAD establishing is the controversial role of transcription. Several studies have shown that inhibition of transcription by α-amanitin does not affect the formation of TAD structures significantly at ZGA in either mouse or *Drosophila* embryos^19–21^. However, a more recent study showed that TAD establishment in human embryos requires transcription^32^. Because transcription is carried out by RNAPII and morpholino inhibition of translation can be easily achieved, we decided to take a different approach. To examine the role of RNAPII in TAD establishment, we used morpholinos to inhibit the expression of RPB1, an essential component of the RNAPII complex. So far, the effects of inhibiting transcription on embryo development have not been examined in detail in other models. However, our studies showed that *X. tropicalis* embryos are highly sensitive to the loss of RPB1. Embryo development was severely delayed, and embryos could only survive up to early s11. This finding underlines the pivotal role of transcription as the mRNA provider for protein synthesis, which is critical for cell survival. The development of embryos to s10 suggests that the background of RPB1 may still provide minimal transcription for cell survival as well as TAD formation. However, at s9, the first-wave TAD formation was severely disrupted, indicating an important role of RPB1 in TAD establishment. At this time, we lack evidence to show if the delayed TAD establishment was due to a direct down-regulation of the amount of RNAPII complex available for binding to DNA or a reduction in transcriptional activity at genes. Nevertheless, our results provide further evidence supporting that at least a component of RNAPII, and very likely transcription itself, is required for the very early stage of TAD formation. The exact mechanism behind the different effects at s9 and delayed s10 remains elusive at this time.

Chromatin modification, protein and epigenetic modification enrichment, gene density and transcription activity are frequently measured for aggregated TADs, which neglects and conceals the variation in single TAD. Based on the analysis of un-clustered TADs, Fudenberg et al. proposed that cohesin-mediated extrusion occurs at loading sites before being stopped at a pair of convergent CTCF binding sites^24^. According to this model, CTCF and cohesin are not expected to be biasedly enriched at either side of TAD borders. Indeed, results from previous publications based on un-clustered TADs show CTCF and cohesion enrichment was roughly equal at both borders if all TADs were aggregated and analyzed.

However, by clustering TADs according to directionality index, our analysis revealed unexpectedly that for most TADs, CTCF and Rad21 are more enriched on one border than on the other. Accompanying this strikingly biased enrichment, the strength of directionality of chromatin interaction at borders shows a similar pattern. Orientation-biased CTCF binding has been proposed to play a role in initiating cohesin-mediated extrusion, which was inspired by the study of *Pcdh* gene loci^26, 33^. Recent findings from the structure analysis of the cohesin-CTCF complex^54^ also provide an explanation for the orientation biased CTCF and cohesin binding at TAD borders. Together with our observation, these data strongly support a model in which the border plays a much more important role in initiating a cohesin-mediated directional extrusion process anchored at orientation-biased CTCF site (Figure 3M).

Our current work suggests that cohesin complex can be loaded and slide in either direction on chromatin until stopped by CTCF binding at a site with a specific orientation. We think that due to structural constraints, cohesin complex and CTCF can only form a unique structure that allows extrusion to happen only in one direction until a barrier stops it. We also speculate that extrusion could be more robust at the anchor site, where the cohesin complex is highly enriched. It then weakens as more DNA extrudes into TAD, where less cohesin is present at the increasingly distant end, which the other border forms. The other border can be any event that counters the weakened extrusion initiated from the first anchor, for example, a convergent site bound by CTCF and cohesin complex, or transcription, which may also pull chromatin to another direction, and reach a balance. However, whether chromatin extrusion does happen as described in the model that we proposed here will require further experimental evidence to prove or refute.

The previous version of the *X. tropicalis* reference genome (v9.1) ^40^ was recently replaced by v10.0. We fixed the numerous errors found in v9.1 and generated a new high-quality reference genome, which together with v10.0, now serves as a valuable resource for the broad research community using *X. tropicalis* to conduct genetic, genomic, molecular, developmental, and evolutionary studies.

In summary, this work provides a systematic analysis of chromatin folding dynamics during embryogenesis through multiple distinct developmental phases. Our results show that TADs are established in at least two waves, with three-phase changes in the internal structure of TADs roughly corresponding to several consecutive developmental stages. We found CTCF, Rad21, and RPB1 are all required for efficient TAD establishment during embryogenesis but differ in their function. We also revealed that RPB1 also plays an important role in developmental stages. More importantly, our data has shed new light on the mechanism of extrusion mediated by the cohesin complex in establishing the TAD structures. Finally, we generated a high-quality reference genome for *X. tropicalis*. Together, these comprehensive datasets will provide a rich resource for studying genome folding principles and the role of the 3D chromatin architecture in gene expression regulation, which governs cell differentiation and decides cell fate.

## METHODS

**KEY RESOURCE TABLE (submitted as pdf file)**

### CONTACT FOR REAGENT AND RESOURCE SHARING

Further information and requests for reagents and resources should be directed to the Lead Contact, Chunhui Hou.

### EXPERIMENTAL MODEL AND SUBJECT DETAILS

#### Frog Strain

*Xenopus tropicalis* frogs were purchased from Nasco (Fort Atkinson, WI, USA) and bred in an in-house facility. All experiments involving frogs were approved by the Institutional Animal Care and Use Committee at the Southern University of Science and Technology, Shenzhen, China. In vitro fertilization was carried out and developmental stages were determined accordiInsing to Nieuwkoop and Faber ^55^. Harvested embryos, cerebral neurons and hepatocytes were isolated from one-year old adult frog and fixed for Hi-C and ChIP experiments. Morpholinos were injected in embryos at the single cell zygote stage.

### METHOD DETAILS

#### Embryos Collection for Hi-C

*Xenopus tropicalis* embryos were obtained at different developmental stages by artificial fertilization. They were cultured in 0.1x MBS medium (1 × MBS: 88 mM NaCl, 2.4 mM NaHCO3, 1 mM KCl, 0.82mM MgSO4, 0.33 mM Ca(NO3)2, 0.41 mM CaCl2, 10 mM HEPES, pH 7.4) at 25°C.

At the desired stages, embryos were fixed for 40 min in 1.5% formaldehyde. Fixation was stopped by a 10 min incubation in 0.125 M Glycine dissolved in 0.1 × MBS, followed by three washes with 0.1 × MBS. Fixed embryos were frozen at minus 80°C in 1.5 ml microcentrifuge tubes (200 embryos per tube).

#### Morpholino Design and Injection

Morpholino antisense oligonucleotides (MO, GeneTools) to *ctcf*, *rad21*, *rpb1*, and control morpholino were separately injected into 1-cell stage embryos from the animal pole with a dose of 10-40 ng per embryo.

Morpholino antisense oligonucleotides for *ctcf*:

5’ GCTTCCGCCATTTCACTTTCCATTT 3’

Morpholino antisense oligonucleotides for *rad21*:

5’ GTGAGCGTAAAACATTTTTCTCTCT 3’

Morpholino antisense oligonucleotides for *rpb1*:

5’ CATGTTTGCGGTGAGACGAGTAC 3’

Morpholino antisense oligonucleotides for *ctrl*:

5’ CCTCTTACCTCAGTTACAATTTATA 3’

#### Hi-C Library Preparation

The generation of Hi-C libraries with low cell number was optimized according to a previous protocol ^11^. Briefly, 100 to 600 embryos were cross-linked with 1% formaldehyde for 40 min using vacuum infiltration. Isolated embryo nuclei were digested with 80U of DpnII (NEB, R0543L) at 37°C for 5 hr. Restriction fragment overhangs were marked with biotin-labeled nucleotides. After labelling, chromatin fragments in proximity were ligated with 4000U T4 DNA ligase for 6 hours at 16°C. Chromatin was reverse cross-linked, purified and precipitated using ethanol. Biotinylated ligation DNA was sheared to 250-500bp fragments followed by pull-down with MyOne Streptavidin T1 beads (Life technologies, 65602). Immobilized DNA fragments were end-repaired, A-tailed and ligated with adaptors. Fragments were then amplified with Q5 master mix (NEB, M0492L). Hi-C libraries were sequenced on the Illumina HiSeq X10 platform (PE 2×150 bp reads).

#### Western Blotting Analysis

*Xenopus tropicalis* embryos and tissues at indicated stages/ages were collected and homogenized in RIPA solution (ThermoFisher) with a proteinase inhibitor cocktail (MedChemExpress). Lysates were mixed with 2 volume of 1,1,2-Trichlorotrifluoroethane (Macklin) and centrifuged at 4°C, 13,000 g for 15 min. Supernatants were mixed with an equal volume of 2× loading buffer and boiled for 5 min. A total of 10 μg of protein was loaded onto a 10% SDS-PAGE gels, electrophoresed, and transferred to a PVDF membrane (Bio-rad). The membrane was blocked by 5% non-fat milk in 1× TBST (a mixture of Tris-buffered saline and containing 0.1% Tween 20) buffer for 1 hour at room temperature and incubated overnight with the primary antibody at 4°C. Anti-RPB1, anti-CTCF, anti-Rad21, anti-beta-Tubulin, and anti-beta-actin were used at concentrations of 1/3000 in 10ml 1× TBST/100mg BSA. Beta-Tubulin and beta-actin were used as loading controls. After five times of washing with 1 × TBST buffer for 10 min, the membrane was incubated with either anti-rabbit or anti-mouse HRP-conjugated secondary antibody (Abmart) for 2 hours at room temperature. The signal was detected using a chemiluminescent western detection kit (Millipore).

#### ChIP Library Preparation

The ChIP assay was performed as described ^56^. Briefly, 200 to 600 embryos were crosslinked with 1% formaldehyde for 40 min using vacuum infiltration. Chromatin was sheared to an average size of 150 bp using a sonicator (Bioruptor Pico, Diagenode). Sonicated chromatin fragments were immunoprecipitated with 3 μg of anti-Rad21 (Abcam, ab992), anti-POLII (BioLegend, 664906), or anti-CTCF (Active motif, 61311). Chromatin-bound antibodies were recovered with 30μl Protein A/G magnetic beads (Millipore 16-663). After reverse crosslinking, ChIPed DNA was recovered using the MinElute Reaction Cleanup Kit (Qiagen 28206) and amplified with the VAHTS® Universal DNA Library Prep Kit for Illumina V3 (Vazyme ND607). Amplified ChIP libraries were sequenced on the Illumina HiSeq X10 platform.

### QUANTIFICATION AND STATISTICAL ANALYSIS

#### Hi-C Sequence Alignment and QC

All Hi-C datasets were processed by using the Juicer pipeline ^57^ and reads with a mapping quality score of less than 1 were filtered out and discarded. The reads were aligned against the *X.tr* v9.1 reference genome, PacBio contigs and our new reference genome respectively. Replicates were merged by Juicer’s mega.sh script. All contact matrices used for further analysis were KR-normalized with Juicer. VC_SQRT-normalized matrices were used when the KR-normalized matrix was not available.

#### Assembly of Genome

We first assembled PacBio reads into raw contigs. *De novo* assembly of the long reads from SMRT Sequencing was performed using SMRT Link’s HGAP4 application with the default parameter. We then scaffolded these raw contigs into chromosome-scale scaffolds. Hi-C data derived from stage 9 cells was selected as assembly evidence considering its low rate of long-range contact and adequate valid interactions. Mapping, filtering, deduplication, merging replicates and scaffolding of contigs based on Hi-C contact were processed by Juicer and 3d-dna ^42^. We skipped the misjoin detection step due to its high false-positive rate in the contigs derived from PacBio long reads. Instead, we manually refined the genome assembly after scaffolding using Juicebox Assembly Tools (JBAT) ^41^ to correct several obvious errors. Note that our new assembly still has some small-scale errors to be corrected.

We then assigned the chromosome number to each chromosome scale scaffolds after the assembly of raw contigs. Genetic markers^58^ were mapped to chromosome-scale scaffolds to determine their chromosome ID and reorient their directions. MAKER^59^ was used to map the previous annotation of *Xenopus tropicalis* to our new genome assembly.

#### Genome Assembly Statistics

For the comparison between previous assembly and our new assembly, we used MUMmer4 ^60^ (command: “nucmer -t 20 -g 50000 -c 1000 -l 1000 --mum”) to align them. Alignments between the two assemblies were then visualized using R basic graphic package. Locations for centromeres of chromosomes were visually determined based on the Hi-C heatmap. For the profile plot of genome assembly, we generated an agp file based on 3d-dna’s output after completing the genome assembling. The profile plot of the genome assembly is based on the agp file.

#### Insulation, TAD, and TAD Border Calling

To check the contact domain properties of each sample, we also calculated directionality index (DI) and insulation score as defined before ^4, 61^ using a parallel script based on 5kb resolution (using triplet format matrix). DI and insulation scores were both calculated with a block size of 200kb (40 bins).

For the domain analysis, we first annotated the contact domain by using the arrowhead method. This step was processed at 5kb and 10kb resolutions using default parameters and merged (retaining the 5kb domain annotation for any pair of domains annotated in both the 5kb and 10kb annotations). The same domain between two domain sets was judged by using Bedtools ^62^ intersect command with “-f 0.9 -r”. For comparison between domain sets from different experiments, we counted overlapped domain by using Bedtools intersect command with “-f 0.7 -r”. We computed the arrowhead corner score of the random control domain set, which was generated by using Bedtools shuffle command for distribution analysis of domain score.

#### Clustering of TADs

We used the K-means clustering method to classify domains from each sample by using deepTools ^63^. Domains were clustered based on the DI values within 10 bins around 5’ and 3’ TAD borders (+/-5bins, 5kb per bin), respectively.

#### TAD Aggregation and Comparison

To further check the validation of the domain, we aggregated each domain set or loop domain set as described before ^29^. After aggregation, we divided each aggregated matrix by their mean value for normalization. Note that for the analysis of the knock-down effect, the aggregated matrices were calculated based on the control domain set.

#### Directionality Bias Index

Directionality Bias Index at 5’ and 3’ borders for each TAD was calculated by the formula below:

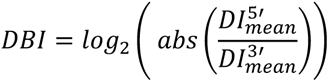

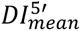: the mean DI value of the 3 bins (15kb) on the right side of the 5’ TAD border.

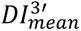: the mean DI value of the 3 bins (15kb) on the left side of the 3’ TAD border.

#### CTCF Motif Enrichment around TAD Borders

To profile the CTCF motif enrichment around domain borders, we first obtained the CTCF motif location and direction in our new genome assembly by using homer scanMotifGenomeWide.pl script ^64^. The enrichment of CTCF in the sense strand and the antisense strand was computed and plotted using deepTools.

#### Compartment Analysis

For the compartment analysis, we calculated eigenvector by using Juicer_tools for each data set at 100kb resolution. H3K4me1 signals from a previous study ^65^ was used to determine the A/B compartment of Hi-C.

To track the dynamics of the domain and compartment structure during embryo development, we performed PCA analysis for both DI value and eigenvector. The R package ggbiplot was used to plot PCA analysis results of DI value and eigenvector.

#### ChIP-seq Analysis

All ChIP-seq reads were mapped to the new genome assembly with BWA ^66^ and analyzed with MACS 2.0 ^67^. All data were normalized against the corresponding input control using the ‘-c’ option of MACS 2.0. Alignments of replicates were merged for downstream analysis.

Signal tracks were calculated by using the ‘bdgcmp’ option of MACS 2.0 with the ‘FE’(fold-enrichment) method. All data for downstream analyses were averaged and extracted from these tracks.

#### Data availability

The sequencing datasets have been deposited under the accession number PRJNA606649.

#### Code Availability

Custom codes used in this study will be provided upon request.

## SUPPLEMENTAL INFORMATION

Supplemental Information includes seven figures and two tables.

## AUTHOR CONTRIBUTIONS

C.H. conceived the study; L.N. and Z.S. performed the experiments; W.S. carried out the data analysis; J.S., J.L. and Y.H. helped with Hi-C libraries preparation; Y.Z., C.F. and P.Z. helped to prepare ChIP-seq libraries; N.H., J.W. and W.W. contributed to Hi-C analysis; C.H. and Y.C. supervised the experiments; C.H. and L.L. supervised the data analysis; C.H. and E.C. wrote the manuscript with input from all authors.

## ACKNOWLEDGMENTS

We gratefully acknowledge financial support from the National Key R&D Program of China (2018YFC1004500), the National Key Basic Research Program of China (2015CB942800), the National Natural Science Foundation of China (31571347 to C.H., 31771430 to L. L, 31671519 to Y.C., and 31701269 to Z.S.), the Shenzhen Science and Technology Innovation Commission (JCYJ20170412152835439), the University of Macau (MYRG 2018-00033-FHS) to E.C. and Huazhong Agricultural University Scientific and Technological Self-innovation Foundation (to L.L.).

## COMPETING INTERESTS

The authors declare no competing interests.

---

**Figure S1.**
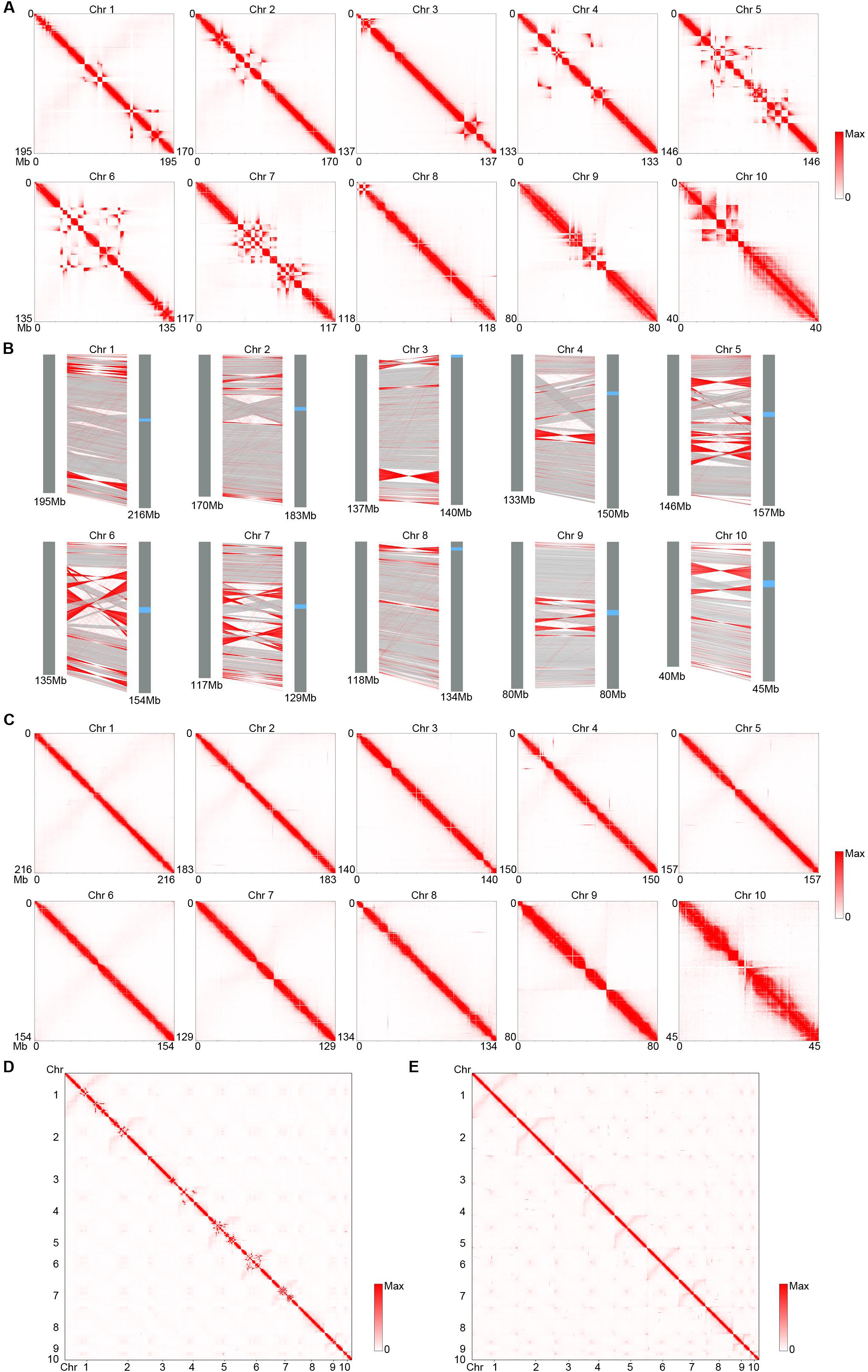
De Novo Assembly of the *X. tropicalis* Reference Genome Assisted by Hi-C and Single-Molecule Sequencing. (A) Heatmap of contact frequency for each chromosome showing the assembly errors using the v9.1 reference genome of *X. tropicalis*. (B) Comparison between v9.1 (left) and de novo assembled chromosomes (right). Red lines show sequences with orientations reversed. (C) Heatmap of each chromosome showing assembly errors are mostly corrected in the new version of the reference genome. (D) Genome-wide contact heatmap showing the lack of centromere interactions using the v9.1 *X. tropicalis* reference genome. (E) Genome-wide contact heatmap showing intra-chromosome arm interactions and inter-chromosome interactions between centromeres.

**Figure S2.**
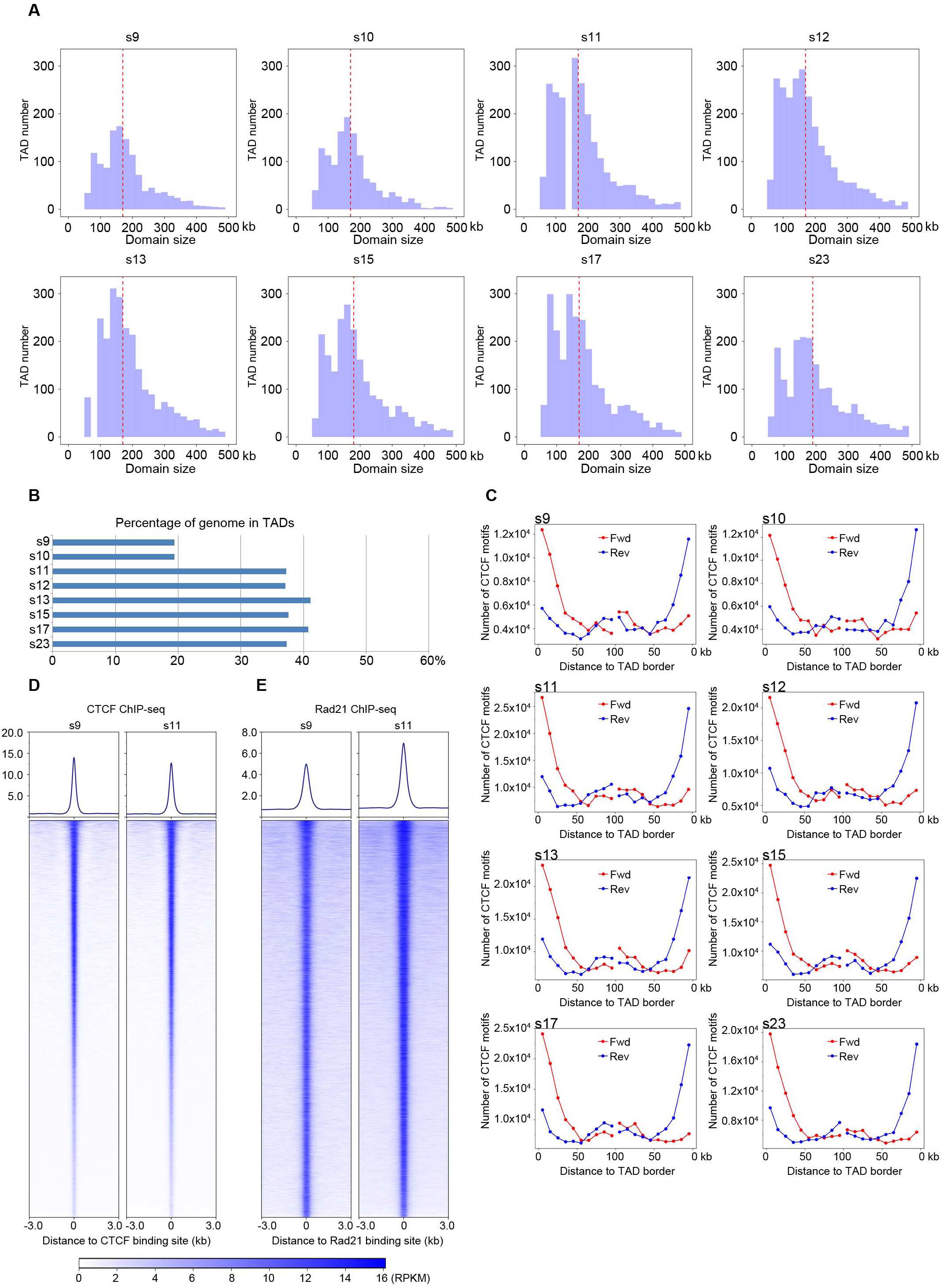
TAD Establishment during the Embryo Development of *X. tropicalis*. (A) Size distribution for TADs identified at different developmental stages. (B) Percentage of genome folded into TAD structures at different developmental stages. (C) Convergent CTCF sites observed at borders of TADs identified at all developmental stages. (D) CTCF ChIP-seq at stage 9 and 11. (E) Rad21 ChIP-seq at stage 9 and 11.

**Figure S3.**
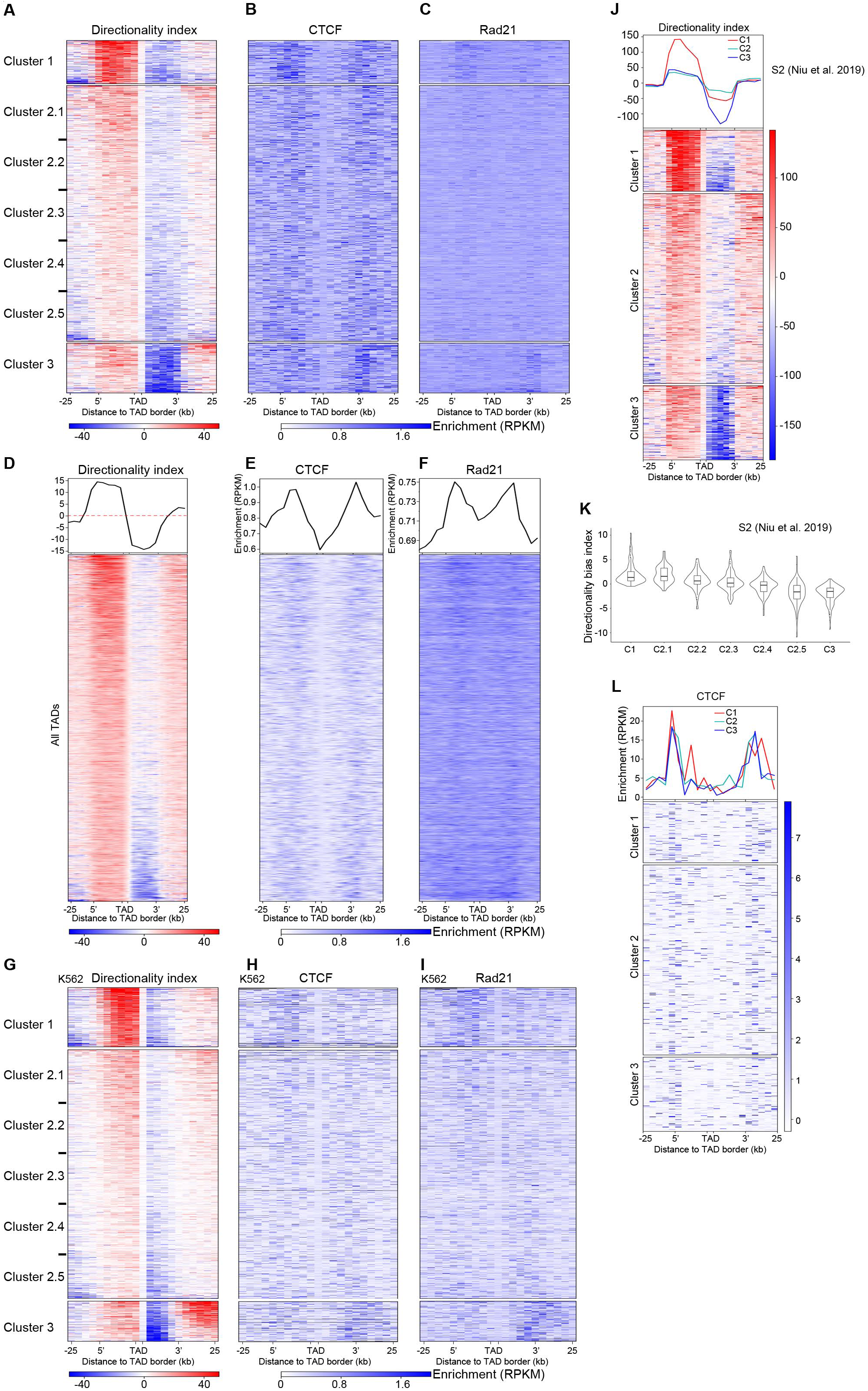
Orientation-Biased CTCF and Rad21 Enrichment at TAD Borders of Higher Directionality Index Values. (A) TADs from stage 13 embryos clustered on directionality index. Cluster 2 is further divided into five sub-clusters with an equal number of TADs. (B) Fold enrichment of CTCF across TAD borders in each cluster. (C) Fold enrichment of Rad21 across TAD borders in each cluster. (D) Average directionality index across borders of all TADs without clustering. TADs are arranged with DI of decreasing absolute value on the 5’ border and DI of increasing absolute value on the 3’ border. (E) Relative enrichment of CTCF across borders of all TADs showing undistinguishable difference in signal strength at borders on both sides of TADs. (F) Relative enrichment of Rad21 across borders of all TAD showing undistinguishable difference in signal strength at borders on both sides of TADs. (G) Directionality index across borders of clustered TADs of human K562 cells. Cluster 2 is further divided into five sub-clusters with same number of TADs. (H) Fold enrichment of CTCF across TAD borders in each cluster of human K562. (I) Fold enrichment of Rad21 across TAD borders in each cluster of human K562. (J) Directionality index across borders of clustered TADs of *Drosophila* S2 cells. (K) Violin plot showing the distribution of directionality bias index of clusters 1, 3 and five sub-clusters of cluster 2 in *Drosophila* S2 cells. (L) Relative enrichment of CTCF at the borders of three clusters of TADs showing the lack of correlation with the inequality in directionality for three clusters of TADs in *Drosophila* S2 cells.

**Figure S4.**
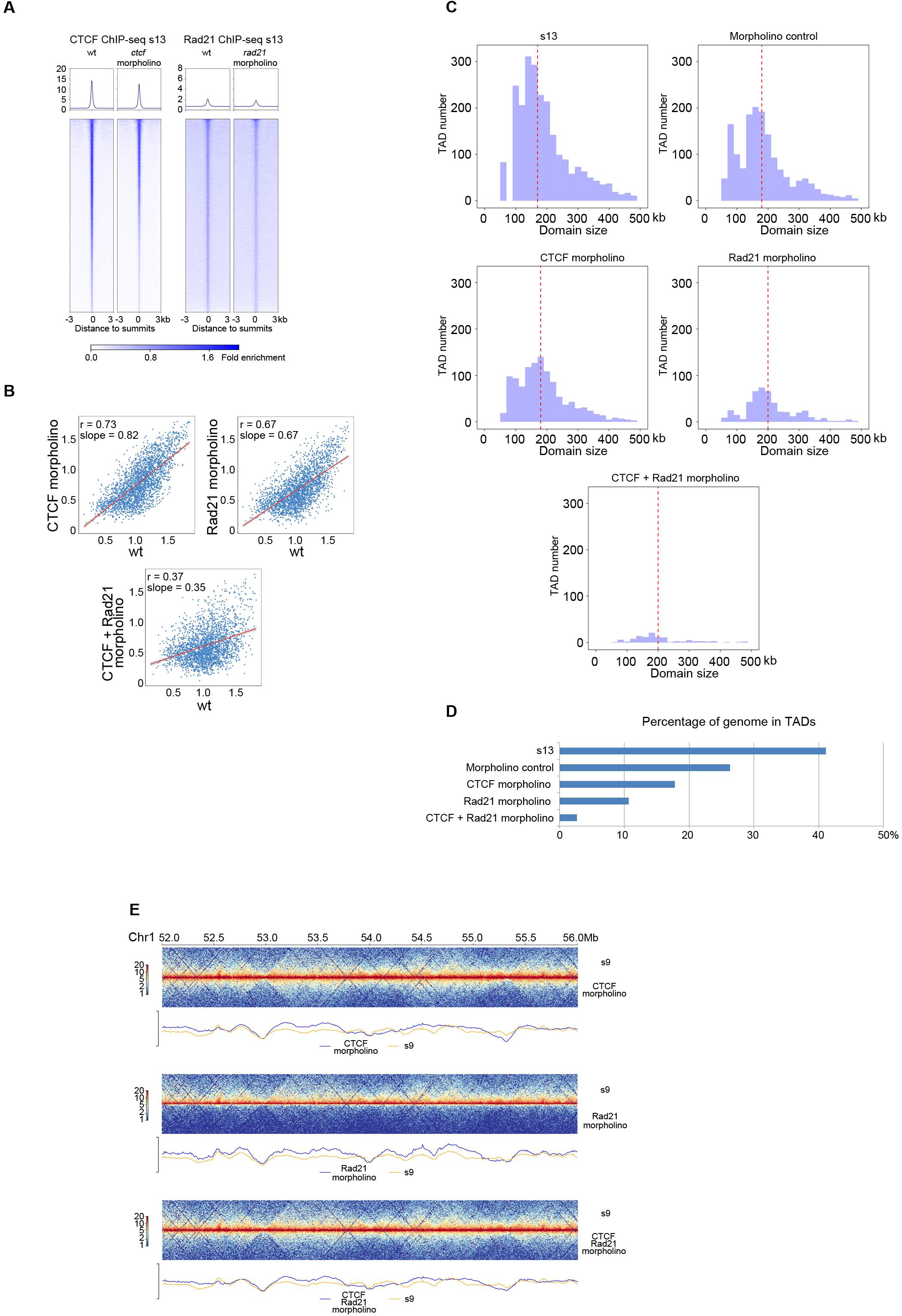
CTCF and Rad21 Knock-Down Effects on TAD Formation. (A) CTCF binding in s13 embryos treated with morpholinos against CTCF. Rad21 binding in s13 embryos treated with morpholinos against Rad21. (B) Scattering plots of insulation scores for the borders of identified TADs in wild type embryos at s13 vs morpholino CTCF, Rad21 and CTCF+Rad21 treated embryos at s13. (C) Number of TADs and the size distribution of TADs. (D) The percentage of genome folded into TADs. (E) Example region to show the knock-down effect on TAD structures.

**Figure S5.**
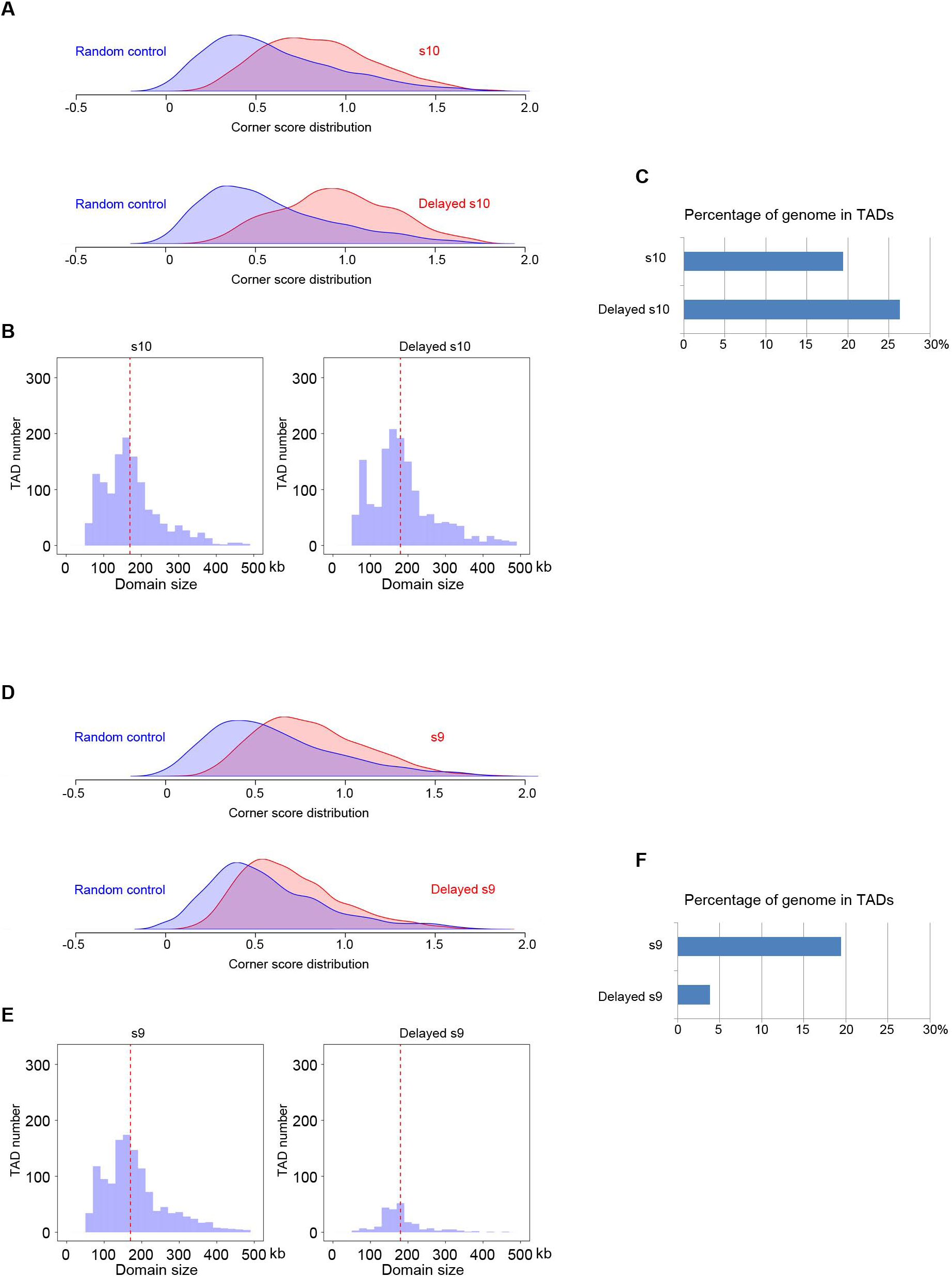
RPB1 Knock-Down Effects on TAD Formation. (A) Arrowhead corner score distribution for wild type stage 10 and RPB1 knock-down embryos at delayed stage 10, respectively. (B) Number of TADs and the size distribution of TADs. (C) The percentage of genome folded into TADs. (D) Arrowhead corner score distribution for wild type stage 9 and RPB1 knock-down embryos at delayed stage 9, respectively. (E) Number of TADs and the size distribution of TADs. (F) The percentage of genome folded into TADs.

**Figure S6.**
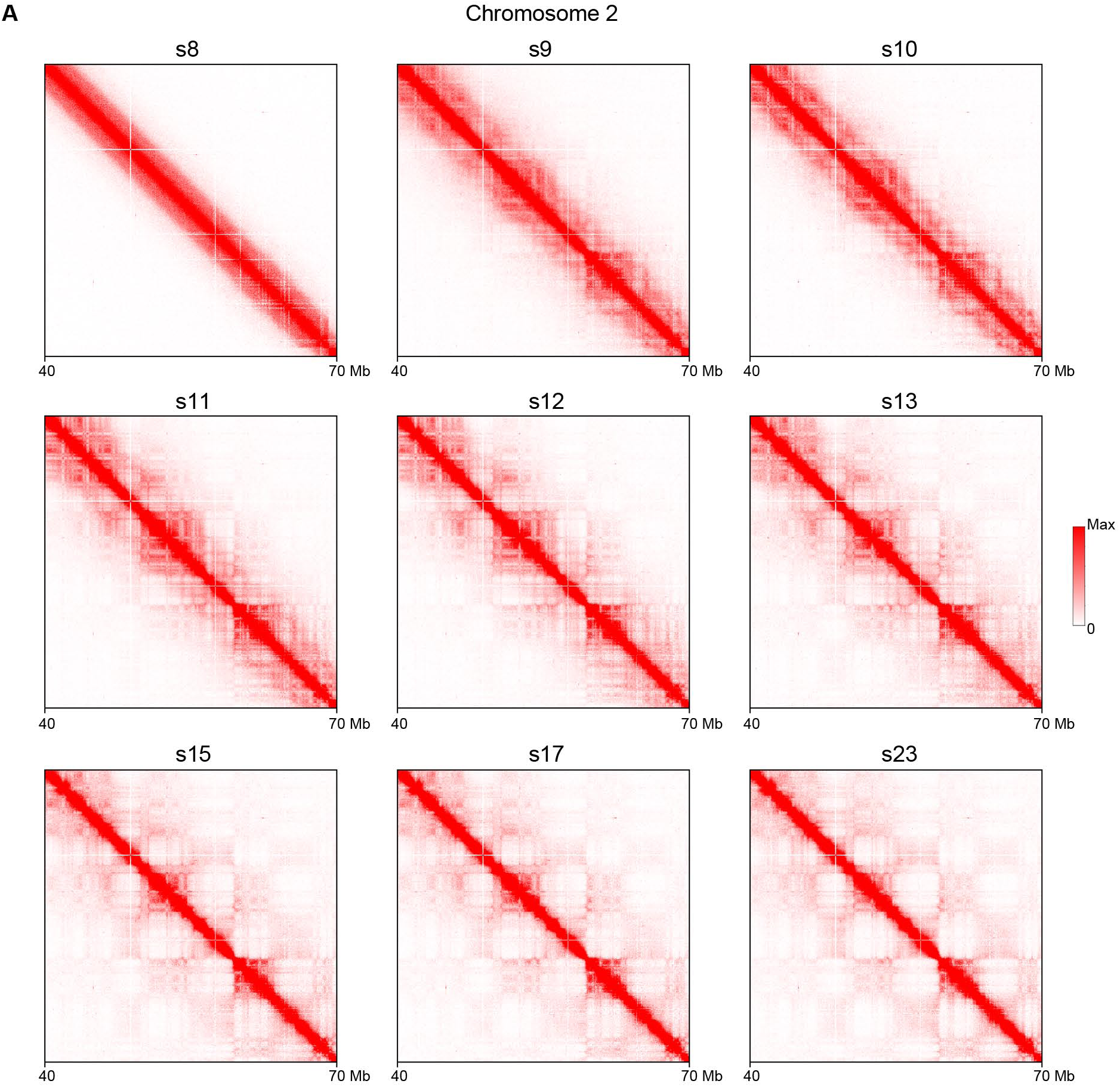
Continuous Compartmentalization of Genome during *Xenopus tropicalis* Embryo Development. (A) Chromatin contact heatmaps at 50kb resolution for a 30Mb region in chromosome 2 from multiple developmental stages showing continuous compartmentalization.

**Figure S7.**
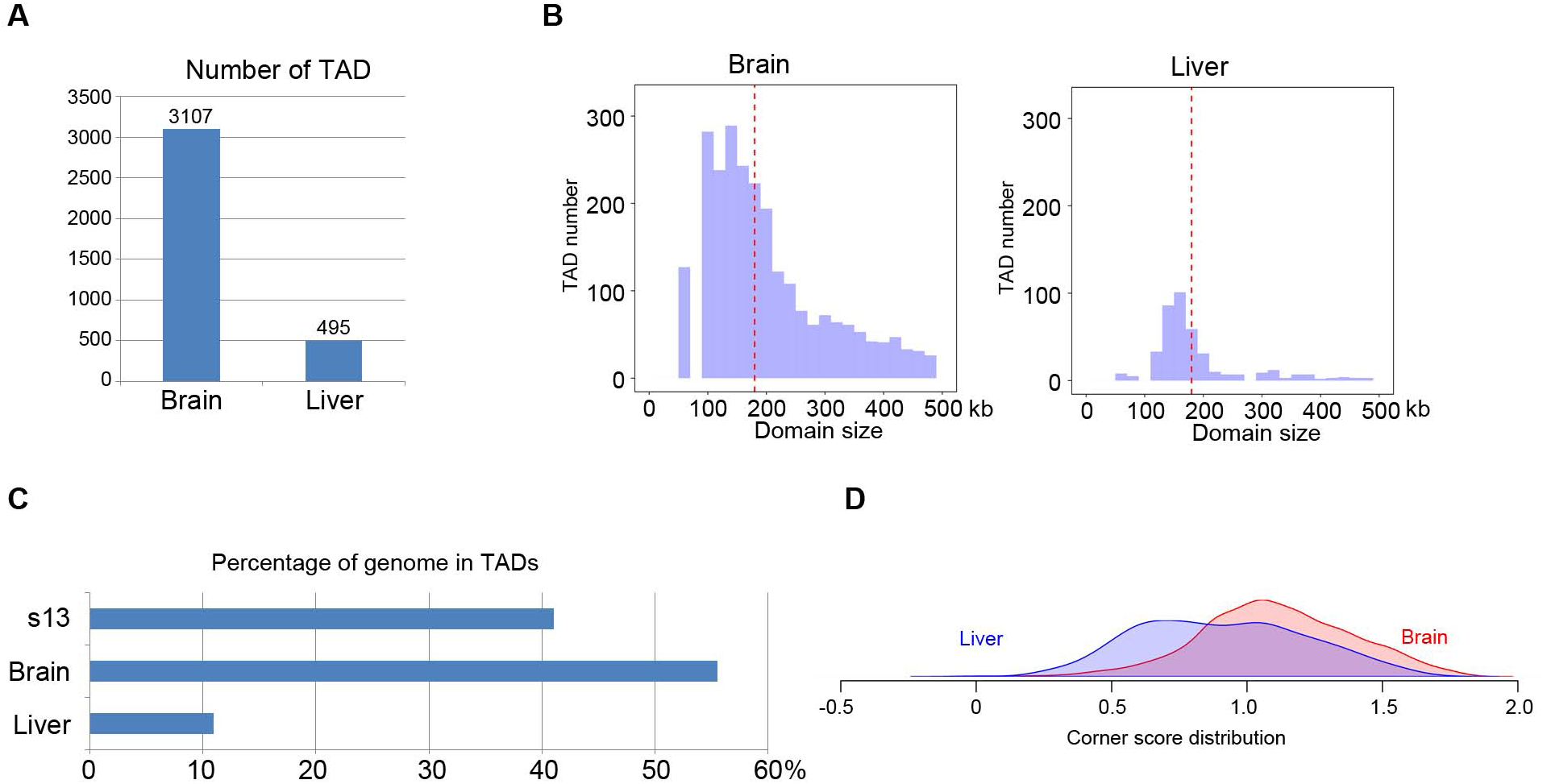
TAD Structure in Terminally Differentiated Brain and Liver Cells. (A) Number of TADs in brain and liver cells. (B) Size distribution of TADs in brain and liver cells. (C) The percentage of genome folded into TADs in brain and liver cells. (D) Comparison of arrowhead corner score distribution between brain and liver cells.

**Table S1.**
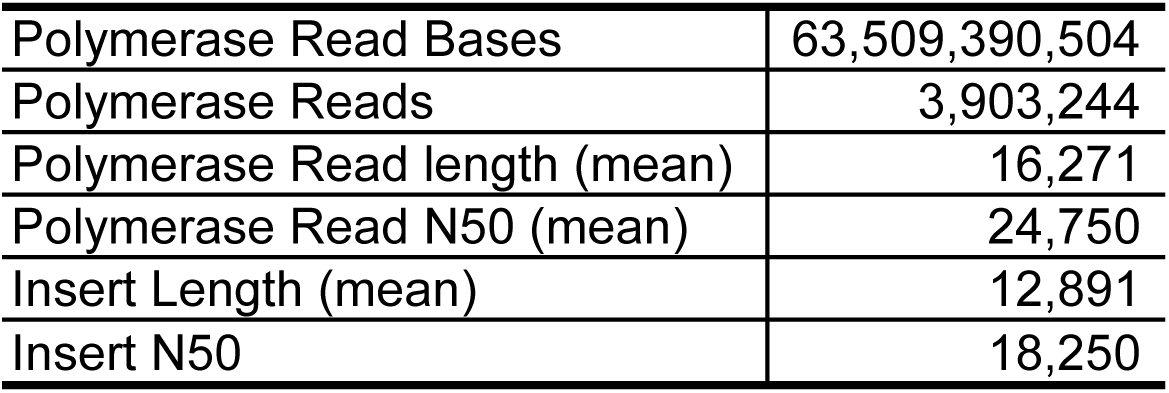
PacBio Sequencing Statistics

|  |  |
| --- | --- |
| Polymerase Read Bases | 63,509,390,504 |
| Polymerase Reads | 3,903,244 |
| Polymerase Read length (mean) | 16,271 |
| Polymerase Read N50 (mean) | 24,750 |
| Insert Length (mean) | 12,891 |
| Insert N50 | 18,250 |

**Table S2.** Illumina Paired-End Sequencing Statistics of Hi-C Libraries

| Experiment | Sequenced Read Pairs | Alignable<br>(Normal+Chimeric Paired) | Hi-C Contacts | Resolution<br>(bp) |
| --- | --- | --- | --- | --- |
| Stage 8 | 976,673,356 | 745,898,363 (76.37%) | 257,081,734 (26.32% / 59.86%) | 7,000 |
| Stage 9 | 784,636,123 | 574,120,502 (73.17%) | 243,499,872 (31.03% / 63.32%) | 6,850 |
| Stage 10 | 608,770,227 | 452,594,668 (74.35%) | 222,209,130 (36.50% / 66.08%) | 7,500 |
| Stage 11 | 795,496,956 | 595,735,571 (74.89%) | 268,305,965 (33.73% / 64.17%) | 6,450 |
| Stage 12 | 618,193,938 | 467,879,429 (75.68%) | 242,140,555 (39.17% / 66.17%) | 7,000 |
| Stage 13 | 573,907,017 | 436,901,789 (76.13%) | 225,420,739 (39.28% / 66.46%) | 7,550 |
| Stage 15 | 647,718,979 | 489,190,955 (75.53%) | 186,117,573 (28.73% / 59.64%) | 8,350 |
| Stage 17 | 639,147,144 | 475,286,350 (74.36%) | 182,443,225 (28.54% / 60.03%) | 8,550 |
| Stage 23 | 616,716,675 | 462,283,925 (74.96%) | 179,402,834 (29.09% / 59.01%) | 8,500 |
| Stage 9: *ctcf* morpholino | 420,981,292 | 334,322,291 (79.41%) | 167,112,043 (39.70% / 67.26%) | 10,250 |
| Stage 9: *rad21* morpholino | 225,985,793 | 134,896,242 (59.69%) | 52,474,861 (23.22% / 63.49%) | 26,500 |
| Stage 9: *ctcf* and *rad 21* m | 419,469,383 | 318,414,579 (75.91%) | 153,455,403 (36.58% / 66.60%) | 11,000 |
| Stage 13: control morpholin | 382,288,544 | 285,758,614 (74.75%) | 140,551,223 (36.77% / 64.28%) | 11,000 |
| Stage 13: *ctcf* morpholino | 473,927,848 | 367,416,074 (77.53%) | 190,531,912 (40.20% / 66.50%) | 9,150 |
| Stage 13: *rad21* morpholin | 427,093,246 | 321,318,596 (75.23%) | 159,284,248 (37.29% / 64.86%) | 9,850 |
| Stage 13: *ctcf* and *rad21* n | 421,568,407 | 319,998,116 (75.91%) | 158,420,927 (37.58% / 66.06%) | 9,800 |
| Delayed stage 9: *rpb1* mor | 360,731,826 | 267,406,200 (74.13%) | 124,142,617 (34.41% / 63.21%) | 11,750 |
| Delayed stage 10: *rpb1* mo | 404,118,645 | 310,383,621 (76.81%) | 149,876,297 (37.09% / 64.95%) | 10,750 |
| Brain | 840,578,973 | 644,452,794 (76.67%) | 297,993,329 (35.45% / 62.19%) | 5,700 |
| Liver | 923,089,205 | 755,718,430 (81.87%) | 214,650,838 (23.25% / 40.87%) | 6,850 |
| Sperm | 476,688,850 | 369,542,331 (77.52%) | 206,197,477 (43.26% / 59.24%) | 7,250 |

## KEY RESOURCES TABLE

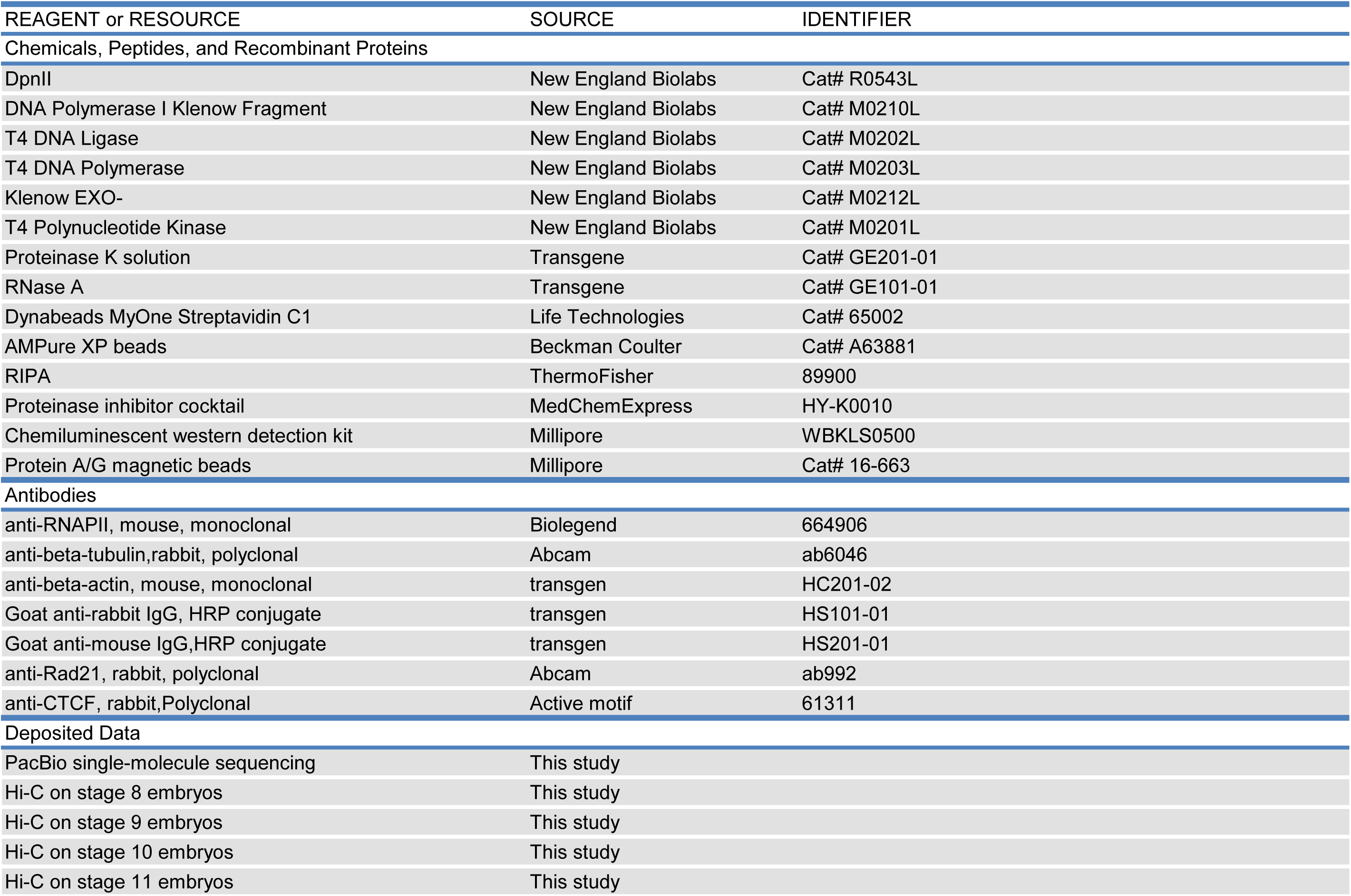
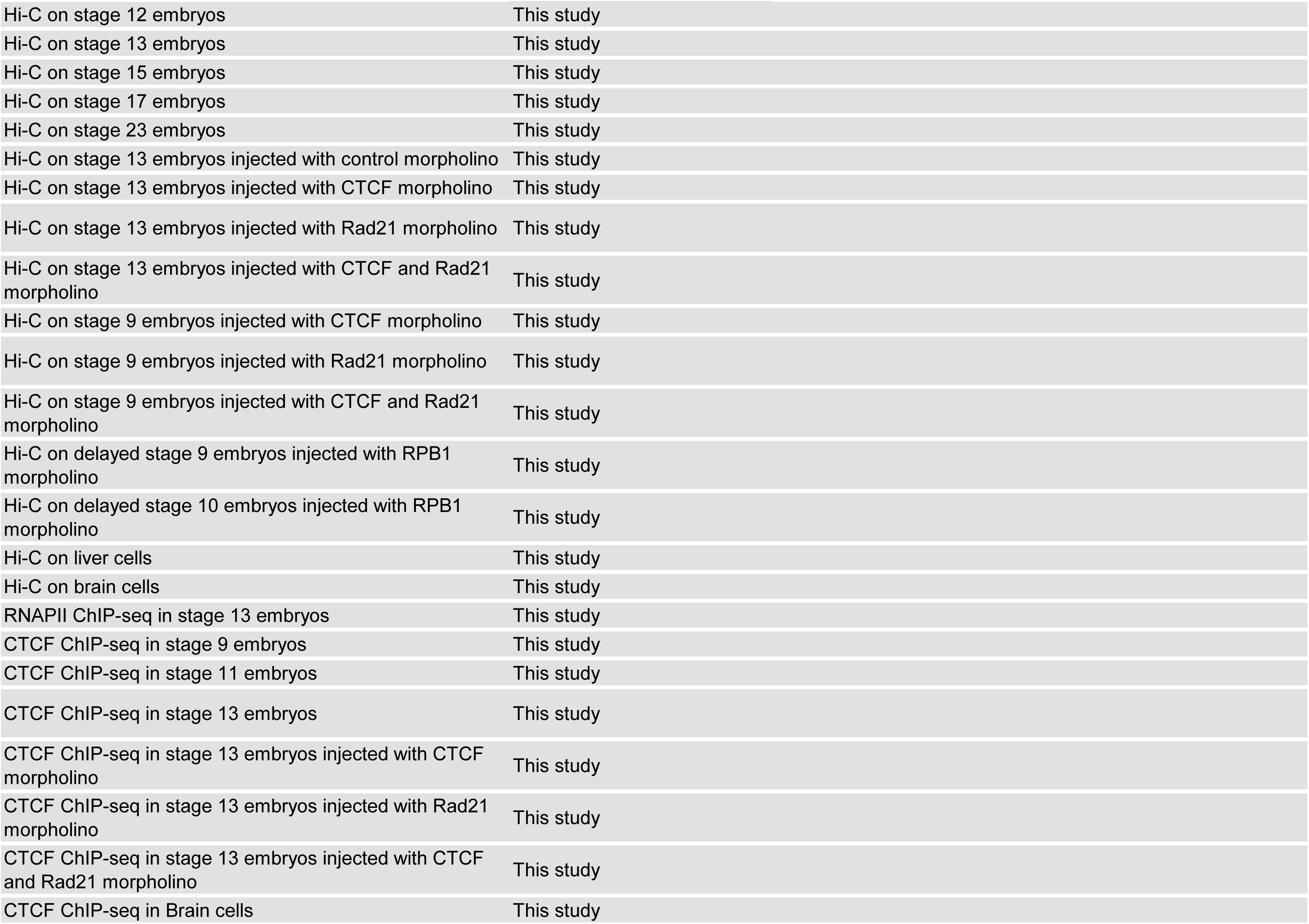
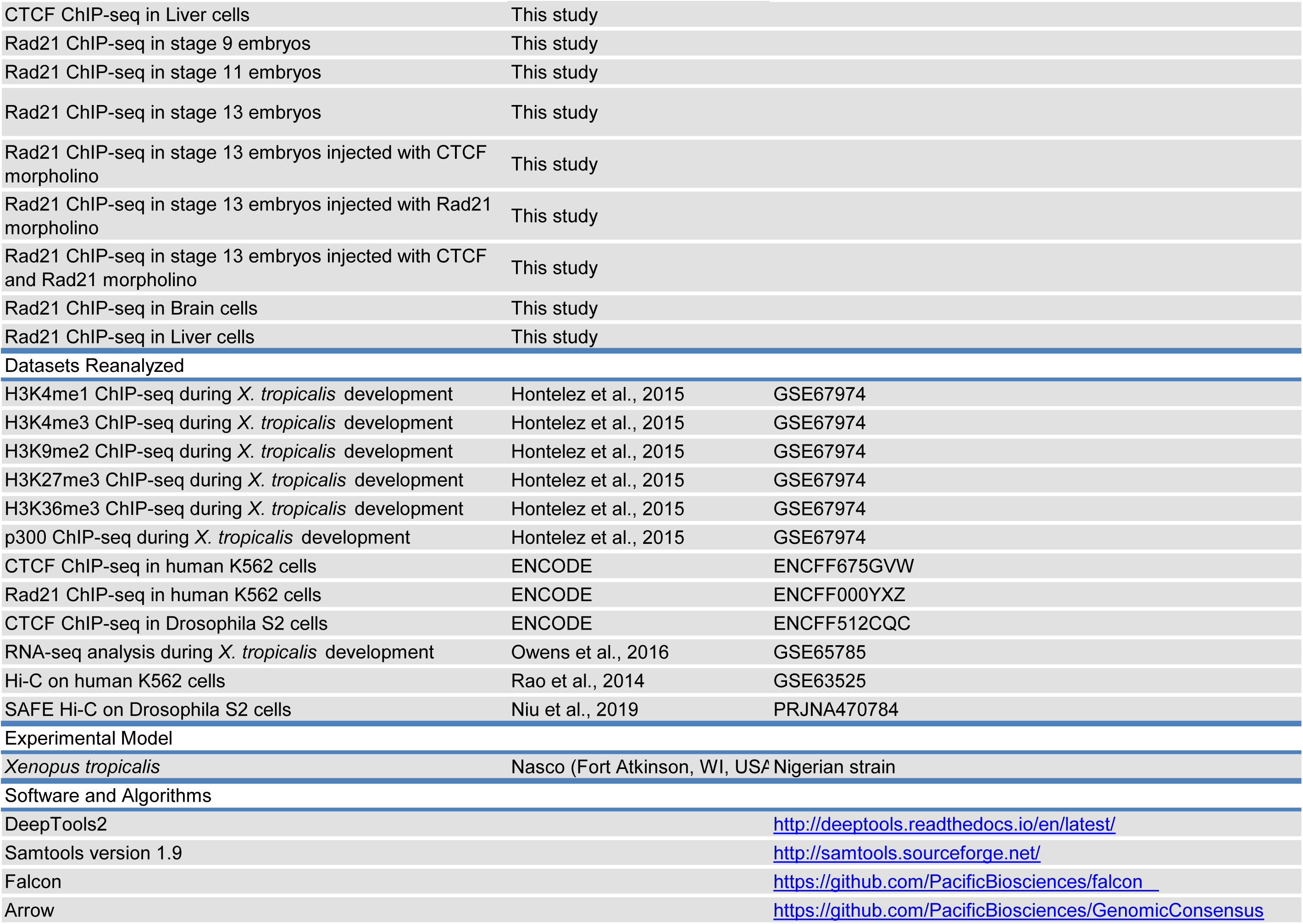
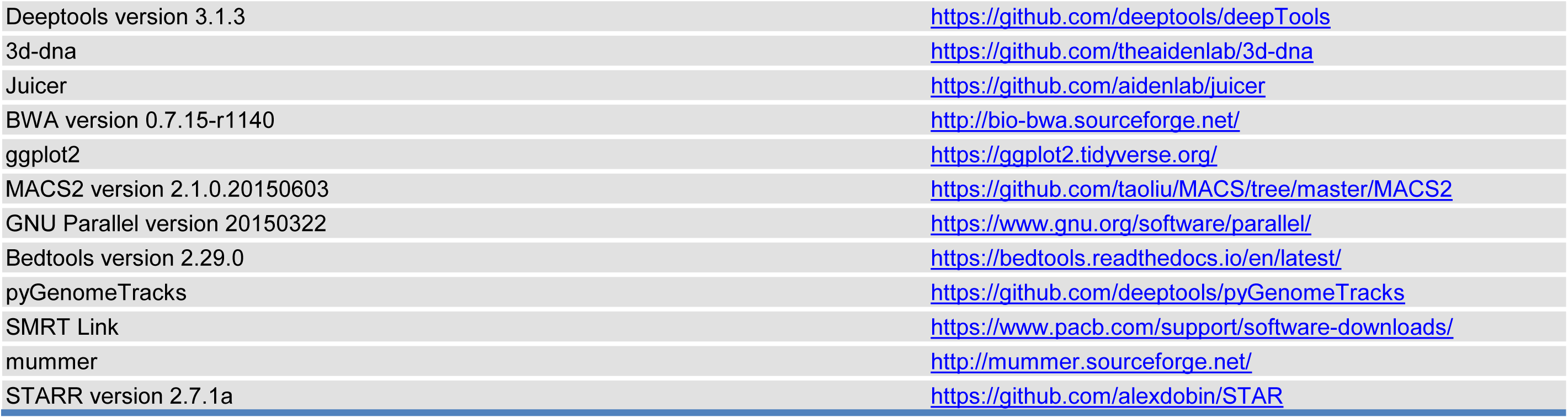

